# Differences in LC integrity and fMRI activations in healthy aging and MCI associated with successful memory encoding

**DOI:** 10.1101/2025.08.29.672869

**Authors:** Lucia Penalba- Sánchez, Yeo-Jin Yi, Elif Kurt, Grazia Daniela Femminella, Clare Loane, Millie Duckett, Nikolaus Weiskopf, Martina F Callaghan, Ray Dolan, Robert J Howard, Emrah Düzel, Dorothea Hämmerer

## Abstract

**INTRODUCTION:** We examined whether the decline of delayed episodic memory in old age and MCI is related to structural integrity of the locus coeruleus (LC). We also tested whether LC integrity was associated with encoding-related brain activity, controlling for emotional salience.

**METHODS:** Participants (28 young adults, 28 older adults, 25 MCI) memorized emotional and neutral images during fMRI. Delayed memory was tested four hours later and compared to immediate memory. LC integrity was measured with a neuromelanin-sensitive sequence.

**RESULTS:** MCI showed overall memory decline, independent of delay or emotional valence, and reduced LC integrity compared to older adults. Across participants, LC integrity correlated with delayed memory, but did not explain performance within OA or MCI. LC integrity was associated with lower activity related to encoding success and emotional salience. Groups remembered emotional better than neutral images, though memory was greatly impaired in MCI, matching with their reduce LC integrity.

**DISCUSSION:** LC integrity was associated with lower brain activity during encoding and emotional salience processing, no clear relationship with delayed memory performance was observed in MCI.

## 1. BACKGROUND

Memory loss is the earliest and most common symptom of Alzheimer’s disease (AD) and primarily associated with a decline in structure and function of the hippocampus and associated areas in memory encoding [1,2].

Similarly affected early on in AD is the locus coeruleus (LC), the primary source of cortical norepinephrine neuromodulation [3]. Its proximity to the fourth ventricle and exposure to small capillaries likely make the LC vulnerable to toxins [4] and early accumulation of tau pathology [5–7]. Based on observations in animal work [8], and inferences from human studies, it is hypothesized that tau protein pathology in the LC in humans also spreads progressively to the entorhinal cortex and other neocortical regions via axonal pathways. This topographically organized progression of tau is a central mechanism in AD, influencing multiple cognitive domains [5] [9] [10].

Early changes in the LC in AD may lead to reduced volume [11] and integrity [12] of the LC, as well as altered noradrenergic neuromodulation [13]. Reduced LC neuron counts, ranging from 30 to 80% in AD and reduced integrity in Mild Cognitive Impairment (MCI) and AD compared to healthy aging [14] are well documented [13] and its association with cognitive outcomes in both populations is receiving increasing attention [7,15,16]. As the LC, through its widespread efferent projections to the hippocampus, helps modulate encoding of, in particular, emotional or salient events into memory, it is assumed to contribute to memory deficits in AD [17] [18]. During exposure to salient or aversive stimuli, LC activity increases, likely as an adaptive mechanism to prioritize the encoding of important experiences [19]. Through its projections to the amygdala [20] and hippocampus [21], the LC modulates emotional memory formation. In line with this, reduced LC integrity is associated with poorer memory particularly for negative emotional stimuli in healthy aging [22].

Despite these findings, the specific functional role of the LC in aging and AD during emotional processing remains unclear. fMRI studies have reported increased amygdala activation in the older adults (OA) in contrast to younger adults (YA) when presented with positive emotional stimuli [23], and reduced amygdala activation along with increased frontal activation when watching negative stimuli [24]. However, whether an impaired LC-NA system in aging, and particularly in individuals with amnestic MCI, is the cause of reduced processing of negative stimuli requires further investigation.

In this study, we present for the first time an LC sensitive task - fMRI paradigm where emotional salience can be assessed independently of cognitive effort in patients with MCI. In addition, we combine this mentioned task fMRI data with structural MRI LC neuromelanin sensitive data that enable to control these two mentioned aspects: salience and effort.

We assess (i) locus coeruleus (LC) integrity, (ii) immediate and delayed episodic memory for emotional and neutral scenes, and (iii) related task-based fMRI activation patterns, focusing on the brainstem and midbrain with an LC-sensitive episodic memory task in young adults (YA), older adults (OA), and individuals with mild cognitive impairment (MCI).

Specifically, we test the following hypotheses: First, we expect increased episodic memory forgetfulness of both neutral end emotional stimuli and reduced benefit from emotional valence in MCI in contrast to the other groups. We expect this to be related to lower LC integrity in MCI. Second, given a more intact LC-NA system we predict stronger activations in regions implicated in negative affect, including the medial prefrontal cortex, anterior cingulate cortex (ACC), LC, amygdala, and insula [33]. Third, we hypothesize higher activation in mid-temporal and frontal regions, hippocampus, and parahippocampus during successful encoding across all groups, with YA showing stronger activations compared to OA and MCI. Finally, we expect increased LC activation in OA compared to MCI during successful encoding, given a more intact LC structure in OA.

## 2. METHODS

### 2.1. Participants

A total of 93 individuals were initially recruited for the study. Data from 11 participants were excluded: 2 older adults (OA) and 4 individuals with mild cognitive impairment (MCI) who did not complete the delayed memory task, and 5 MCI participants whose fMRI data were excluded due to excessive head motion. The final sample consisted of 82 participants: 29 young adults (YA), 28 older adults (OA), and 25 older adults with MCI. Chi-square tests revealed no significant differences in gender distribution across groups (see Table 1).

**Table 1.** Participants demographic, neurologic and cognitive descriptive information.

| Group | YA | OA | MCI | ANOVA *P* |
| --- | --- | --- | --- | --- |
| n | 29(17) | 28(19) | 25(10) | $\chi^2$ ns |
| (males) |  |  |  |  |
| Age | 23.1 (2.7) | 70.6 (6.39) | 72.6 (7.05) | <.001(*f* |
| (SD) |  |  |  | 687)<sup>ab</sup> |
| MMSE | 23.10 (2.8) | 29.48 (0.69) | 25.49 (2.68) | <.001 (*f* |
| (SD) |  |  |  | 44.06)<sup>bc</sup> |
| GMV | 1,180,181.37 | 1,117,230 | 1,064,780.88 | .06 (*f* 2.77)<sup>b</sup> |
| (cm<sup>3</sup>) | (197059) | (200685.47) | (149937.10) |  |
| (SD) |  |  |  |  |
| LC int | 0.13 (0.03) | 0.16 (0.04) | 0.10 (0.03) | <.001(14.38)<sup>d</sup> |
| (SD) |  |  |  |  |

Note. Columns two, three and four display results as mean (standard deviation). Column five shows the Chi squared for age and ANOVA *p* values. *n* = sample size; MMSE = Mini Mental State Examination; GMV = Gray Matter Volume; LC int = Locus Coeruleus Integrity of the 3 voxels with maximum integrity (for LC int score see Methods); YA = Younger Adults; OA = Older Adults; MCI = Mild Cognitive Impairment). a) Significant differences between YA and OA, b) Significant differences between YA and MCI, c) Significant differences between OA and MCI, d) Significant differences between all three groups.

Exclusion criteria included any self-reported psychiatric or neurological disorders. All MCI participants had received a formal diagnosis from a clinician and had a Mini- Mental State Examination (MMSE) [26] score between 22 and 27 (see Table 1).

YA were recruited via email from a participant database maintained by the Institute of Cognitive Neuroscience at University College London (UCL). Older adults were recruited through advertisements, flyers, or via clinical teams when attending memory clinics alongside friends or relatives with MCI. MCI patients were recruited through the Join Dementia Research database and from local memory clinics within the Camden and Islington NHS Foundation Trust and the North East London NHS Foundation Trust. Written informed consent was obtained from all participants. The study was conducted in accordance with the Declaration of Helsinki and received ethical approval from the University College London Research Ethics Committee (ethics reference number: 17/0091).

### 2.2. Study materials and paradigm

The task was programmed using MATLAB R2015a (version 8.5.0.197613, The MathWorks, Inc, 2015) and the Cogent 2000 software. Scene images with both emotional and neutral content were drawn from a validated set taken from the International Affective Picture System (IAPS) database[27] Each scene image belonged to one of four groups, each containing 22 stimuli: emotional outdoor, emotional indoor, neutral outdoor, or neutral indoor. All stimuli were adjusted to 50% luminance to control for brightness effects on pupil dilation, which was recorded during the experiment; these results are not reported in this manuscript. Binary checkerboard noise patterns at 50% luminance served as background stimuli. Event-related fMRI was recorded for the incidental encoding memory task (Figure 1A), followed by an immediate recognition memory test 30 min after encoding and a delayed recognition memory test 4 hours after encoding (Figure 1B). Participants were asked to classify stimuli as indoor or outdoor to ensure sufficient attentional processing during the incidental encoding task. To ensure sufficient trial numbers for the analysis of subsequent memory effects[28], and given the expected memory decline and attentional strain in the MCI group, we combined trials correctly identified as ‘old’ in either the immediate or delayed recognition test into a single ‘remembered’ condition.

**FIGURE 1.**
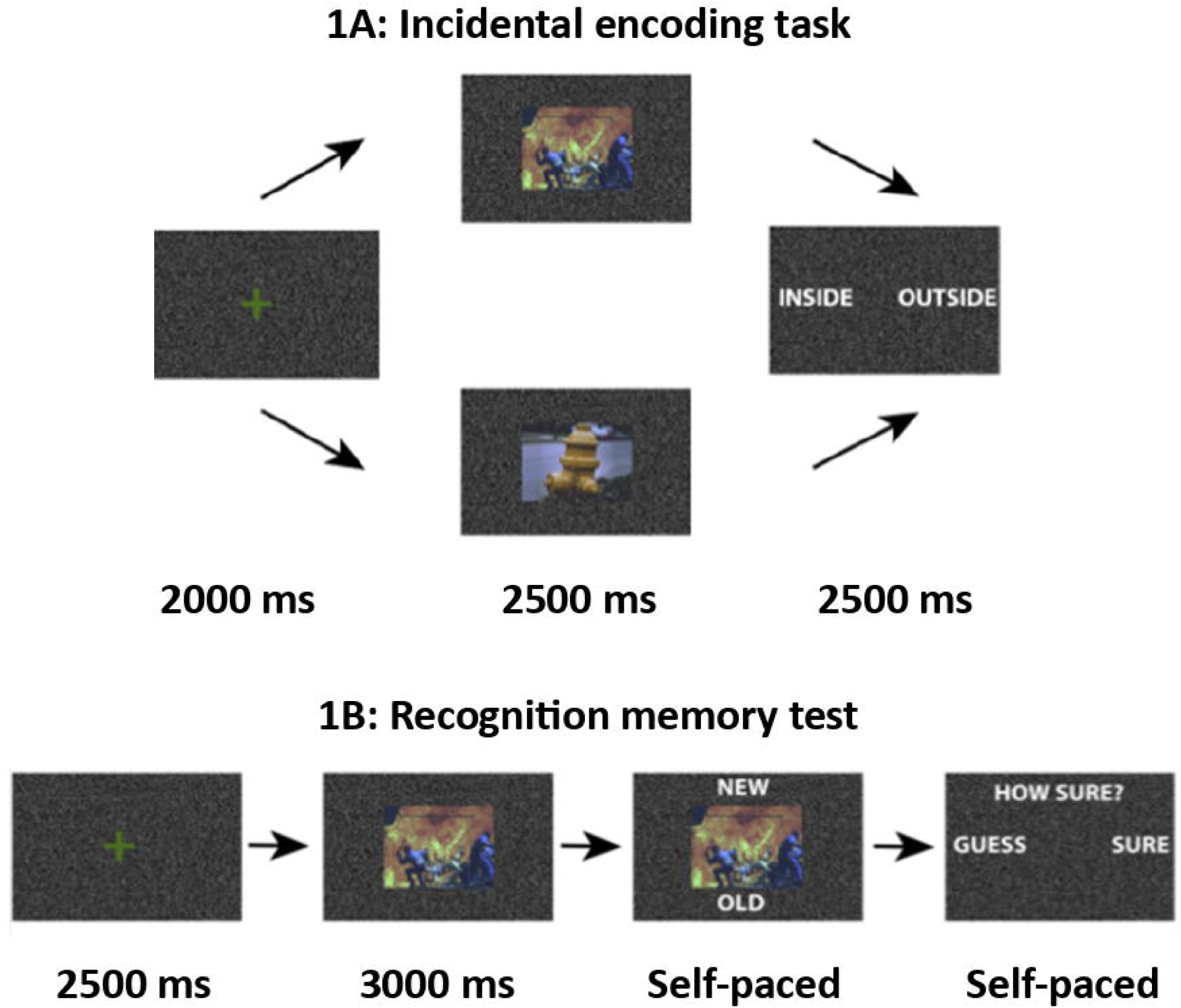
**(A)** Conducted inside the scanner: Participants view an image of a scene, which may contain either negative or neutral affective content. As a cover task to ensure attentional processing of the scene, participants are instructed to press the right mouse button if the scene takes place outdoors (e.g., in a park or on a street) and the left mouse button if it takes place indoors (e.g., at home or in a restaurant). **(B)** Conducted outside the scanner 30 min after encoding (immediate memory) as well as four hours later (delayed memory) (n = 44 images previously shown in the scan and n= 44 new images). Participants view an image and indicate whether it was previously shown during the scanning session ("old") or if it is a novel image ("new"). Additionally, they specify their confidence level by indicating whether they "guess" or are "sure” as well as evaluate the emotionality of the scence as “very emotional” or “not very emotional”. Half of the trials displayed are emotional and half are neutral (n = 22 new emotional, n = 22 new neutral).

**FIGURE 2.**
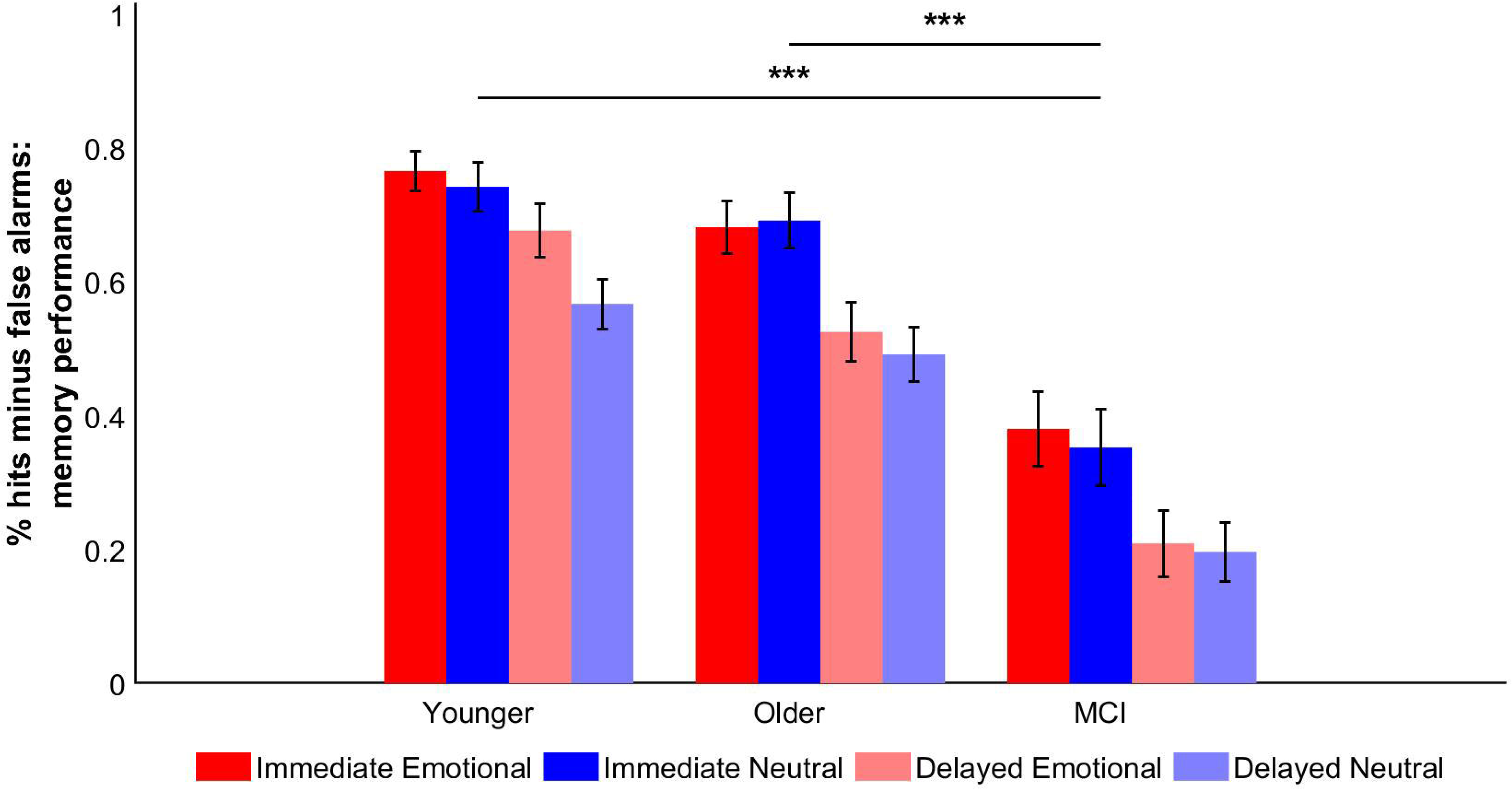
shows mean memory performance (% hits minus % false alarms) of the immediate (in darker colour) and the delayed memory tests (in lighter colour) for each group and each type of image (negative emotional in red and neutral in blue). Lines with asterisks indicate the main effect of group in memory performance (MCI < younger adults and older adults).

During the incidental encoding task in the scanner, 88 trials were presented: 44 emotional and 44 neutral. Scenes were presented for 2.5 seconds, preceded by a fixation cross which varied between 1.5 and 5.5 seconds (mean duration 2 seconds), followed by the outdoor-indoor question which was presented for 2.5 seconds. The total task duration was approximately 10 minutes. In the delayed memory test, 88 trials were presented, comprising 22 new emotional images, 22 new neutral images, 22 emotional images previously shown during the scan session, 22 neutral images previously shown during the scan session. Moreover, half of the scene stimuli were indoor and half outdoor. Scenes were presented for 3 seconds, preceded by a fixation cross which varied between 2.25 and 2.75 seconds (mean duration 2.5 seconds), followed by the old-new question and guess-sure question which were presented until a response was given. Total task duration was approximately 12 minutes, depending on reaction times.

### 2.3. Imaging parameters

All MRI scans were acquired on a Siemens MAGNETOM Prisma (Siemens Healthcare, Erlangen, Germany) equipped with a 64-channel head coil. Participants lay supine, with foam padding to minimize head movement. Sequences included both structural and functional acquisitions, as well as additional scans for geometric distortion correction.

#### 2.3.1. Structural MRI acquisition

Whole-brain T1-weighted structural images were acquired using a 3D Magnetization Prepared Rapid Gradient Echo (MPRAGE) sequence to support co-registration of the functional images to structural group templates and evaluate individual volumetric indices (1.0 mm isotropic voxels, TR = 2530 ms, TE = 3.34 ms, TI = 1100 ms, flip angle = 7°, field of view = 256 × 256 × 176 mm^3^). A dual-echo gradient-echo field map (TR = 1020 ms, TE1 = 10.00 ms, TE2 = 12.46 ms, flip angle = 90°, resolution = 3.0 × 3.0 × 2.0 mm) was acquired for B0 inhomogeneity corrections in functional recordings.

In order to assess the LC in the brainstem, a 3D magnetization transfer-weighted (MTw) FLASH sequence (voxel size = 0.6 × 0.6 × 3.0 mm, TR = 24.50 ms, TEs = 3.35/6.15/8.95/11.75 ms, flip angle = 12°, field of view = 186 × 212 × 60 mm) was used to obtain high-resolution images of LC integrity. The acquired MTw images were averaged across 3 repetitions and 4 echoes. It is assumed that neuromelanin deposits in the LC make the LC appear hyperintense in this so-called ‘neuromelanin-sensitive’ contrast [22,29]. As neuromelanin accumulates within the LC cells, a higher signal intensity is interpreted as a higher cell density in a given voxel or higher structural integrity of the LC [30]. Quality assurance checks of acquisition were performed by the radiographer to prevent imaging data containing artefacts (e.g. movement artefacts). The sequence was repeated if an artefact was found during this visual inspection.

#### 2.3.2. Functional MRI acquisition

As the goal of the present study was to explore memory and emotional processing with a focus on the brainstem and midbrain, data was optimised so that the resolution and coverage enabled the assessment of deep and small structures (e.g., LC and amygdala). A high resolution 2D T2*-weighted whole brain multi-band echo-planar imaging (EPI) sequence was acquired (2.0 mm isotropic voxels, TR = 1450 ms, TE = 35 ms, TA = 14:40, flip angle = 70°, multiband factor = 4, no in-plane parallel imaging imaging, Bandwidth = 2246 Hz/Px, field of view = 212 mm × 212 mm × 144 mm, 72 interleaved slices).

#### 2.3.3. fMRI pre-processing steps

fMRI data were preprocessed in combination of SPM12 (Wellcome Centre for Human Neuroimaging, UCL, 2012; http://www.fil.ion.ucl.ac.uk/spm12.html), Advanced Normalization Tools (ANTs; version 2.5.1 post39-g2236a75; http://stnava.github.io/ANTs/; preprocessing steps involving this toolbox are detailed in subsection 2.3.4 of this manuscript), FSL (version 6.0.1; https://fsl.fmrib.ox.ac.uk/fsl/, Analysis Group, FMRIB, Oxford, UK), and Freesurfer (Version 7.1; http://surfer.nmr.mgh.harvard.edu/, Martinos Center for Biomedical Imaging, Charlestown, Massachusetts). DICOMs were converted to NIfTI, and slice timing was corrected using the multi-band parameters from the JSON sidecar of the fMRI sequence. Realignment and unwarping was performed with SPM12 using voxel displacement maps calculated from the double-echo B_0_ field maps, with mostly default settings except for a spatial separation of 2 mm, a 3 mm FWHM realignment kernel, and 4^th^-order interpolation given the higher than usual spatial resolution in acquired data. This step generated a mean functional image, which was later used as a representative image of a fMRI time-series dataset during co-registration. The 4-D fMRI were then smoothed with a 2 mm full-width-at-half-maximum (FWHM) Gaussian kernel using the Smooth function of SPM12. Afterwards, first-level contrasts were estimated in each subject’s native space without registration or normalization. In addition, T1-weighted structural images of all participants were bias-corrected in ANTs (N4BiasFieldCorrection) prior to template generation. Study (a template created from all subjects) as well as three group-specific templates (created including only YAs, OAs, and MCI patients, respectively) were constructed using antsMultivariateTemplateConstruction2 [31] (template construction details available at: https://github.com/alex-yi-writes/LC-SpatialTransformation2021).

#### 2.3.4. Precise spatial transformations of structural and functional LC imaging data

To ensure sufficiently precise spatial transformations of structural and functional imaging data across subjects to examine activations in the small brainstem structures such as the LC, the procedures detailed in Yi et al. (2023)[32] were modified and performed to fit the study objectives and dataset properties. Firstly, individual structural images were nonlinearly transformed to the study-specific and group-specific templates via the antsRegistrationSyN.sh function to facilitate the normalization of mean functional images and the statistical contrast maps onto the template images[31].

Afterwards, mean functional images were rigidly registered to the corresponding structural images, also via antsRegistrationSyN.sh, generating affine transformation matrices. Finally, concatenating the transformations (affine matrices and warp fields) acquired from each step, the statistical contrast maps in their native space were transformed directly into the study- and group-specific space with only one interpolation step using the antsApplyTransforms function of ANTs with linear interpolation setting[31].

For functional LC data, a quality assessment procedure for sufficiently precise spatial transformations proposed by Yi et al. (2023)[32] was subsequently employed. Specifically, six landmarks were manually placed in brainstem regions on individual mean functional images in four group- and study-specific template spaces, i.e. study, YA, OA, and MCI templates. A rater with extensive experience in LC dataset evaluation performed this landmark placement (more detailed procedure of the landmark selection and placement protocols are described in Yi et al. 2023)[32]. As shown in Supplementary Fig. 1, landmark deviations in the LC area were on average 0.79 mm and did not exceed the typical half-width of the LC (2 mm)[33].

For structural LC data, individual LC masks were segmented using ITK-SNAP. The combined region of two independent ratersconsensus(DH and YY) was used to generate the final mask, as previously described [34]. LC integrity was computed as the contrast ratio (CR) of signal intensity in the joint mask across both raters divided by the signal intensity of a reference region in the pons, not assumed to contain neuromelanin [12,35,36]. The size and location of the reference region were based on previous literature[37]. Sørensen–Dice coefficients for inter-rater consistency across the three groups were .70 (YAs), .65 (OAs) and .57 (MCI), respectively. LC volumes and LC integrities were not correlated within groups (YAs: r = .22, p = .25; OAs: r = .19, p = .31; MCI: r = .28, p = .12), a result that is not surprising considering that volume indicates the LC extension while integrity is assumed to reflect neuronal density within the LC, which can decline differentially [13]. Finally, we calculated a contrast ratio per participant, using the mean signal of the three brightest voxels in LC and pons reference masks as a more robust LC integrity measure [35].

#### 2.3.5. Masks of small structures for 2^nd^ level analysis (Small Volume Corrections)

For small regions of interest, small volume corrections were applied during statistical analyses [38,39]. For this, region-specific masks were generated for the group and study templates for the following areas: LC, thalamus, hippocampus and amygdala. The LC masks were derived by transforming a meta LC template published by Dahl et al. (2022)[40] into each template space. For the thalamus, hippocampus, and amygdala, the masks were extracted by transforming the whole-brain parcellation maps obtained from each participants structural T1w images, processed using Free surfer’s recon-all function (version 7.4.1, Dale et al, 1998 [41]).

### 2.4. Behavioural analysis and LC integrity

Memory performance (% hits minus % false alarms) of the encoded scenes, as well as LC integrity values were computed and compared among the YA, OA, and MCI groups using one-way ANOVAs, (*p* threshold of *p* <0.05) followed by Bonferroni post hoc tests for pairwise comparisons in SPSS version 29.

### 2.5. Task fMRI analysis: General linear model

In order to take into account atrophy differences across younger adults, older adults, and MCI patients groups (cf. Table 1) two approaches to spatial transformation involving advanced warping methods (SyN-based algorithms) [31] were employed: (i) transforming images to a study-specific template based on averages of anatomies across all three groups; and (ii) transforming images to separate group-specific templates, based on averages of anatomies within each group. If possible, normalizing all participants into a shared space is preferable as it allows voxel-level contrasts examining also partially activated ROIs between groups [42–45]. However, if for example the MCI group showed substantially smaller or fewer activated regions in the study-specific templates space (compared with its own group-specific template), this might indicate registration distortions due to atrophy differences across groups which affect significances at the group level in a joint template in particular in a more atrophied group. To evaluate this, prior to investigating emotional and memory-related contrasts, a standard “sanity-check” contrast was evaluated comparing the study- specific and group-specific templates to ensure consistency of activation patterns in between study and group specific templates. After confirming that both templates produced comparable results motor related brain regions, e.g., supplementary motor cortex and precentral gyrus (See supplementary figure 2 and supplementary table 1), study-template based analyses were used for the main emotional and memory contrasts. Event-related regressors and contrasts of interest are summarised in Table 2. Additional models explored emotional processing and memory encoding while controlling for potential confounds or covariates (age, grey matter volume, and LC integrity).

**Table 2.** Event related GLMs, regressors and contrasts.

| **Event related GLM** | **Regressors** | **Contrast of interest** |
| --- | --- | --- |
| 1. Motor (sanity check) | Emotional, neutral, indoor, fixation cross, response, remembered. | Response > fixation cross |
| 2. Emotional processing | Emotional, neutral, indoor, fixation cross, response, remembered. | Negative images > Neutral images |
| 3. Memory encoding | Emotional, fixation cross, response, remembered, not remembered. | Remembered images > not remembered images |

Note. Table 2 shows the GLM = event related General Linear Model; Regressors = the covariates regressed in the model; and the Contrasts of interest = the target contrasts.

Functional activations within and between pairs of groups were examined using one- sample and two-sample t-tests in SPM12, respectively. For the sanity check contrast, the binarized study and group template gray matter masks were used to inspect whole brain level activations, while the binarized study template gray matter mask was applied for the emotional > neutral and remembered > not remembered contrasts. This approach excluded non-gray matter tissues (i.e., cerebrospinal fluid or white matter) from the results via an inclusion gray matter mask based on segmentations of the study /group templates. A threshold of p_uncorrected_ < .001 at the voxel level and p_uncorrected_ < .005 cluster level was used [46].

For the more exploratory assessments of significant clusters in cortical and subcortical areas, we employed a two-step cluster-extent thresholding procedure to identify robust activations while balancing sensitivity and specificity. Statistical maps were initially thresholded at *p*_uncorrected_<.001. Afterwards, data-driven cluster extent threshold was defined by identifying a voxelwise False Discovery Rate (FDR) correction at *q*<.005 on the same contrast[47]. The size in voxels of the largest FDR-surviving cluster was noted. This FDR-derived cluster extent threshold (k) was then applied to the initially thresholded map. Only clusters whose spatial extent exceeded this value are reported as statistically significant. This method warrants that the reported clusters are not only spatially coherent but also meet a mathematically objective criterion for significance derived from the FDR framework.

For confirmatory small volume corrected (SVC) analyses, of the following regions: thalamus, amygdala, LC, and putamen, significance assessments were corrected for multiple comparisons using family wise error correction (FWE), a more conservative measure to assessing the ratio of falsely rejected tests to all tests performed, with a voxel threshold of p <0.01 as these regions are prone to present a lower signal to noise ratio. Findings reported as a trend were significant with a *p*-value between 0.05 and 0.1.

## 3. RESULTS

### 3.1. Behavioural results

Behavioural results (Table 3 and Figure 1) examine the recognition memory performance as operationalised by Hits – False Alarms considering the type of image memorised (neutral or emotional) and the time (immediate memory recognition and delayed memory recognition).

**Table 3.** Behavioural results: within-Subjects Effects and group interaction effects.

| Effect | df | F | *p* | $\eta^2p$ (Effect Size) |
| --- | --- | --- | --- | --- |
| Type image (Neutral vs. Emotional) | 1, 79 | 5.46 | .022 | .067 |
| Type image × group | 2,79 | 1.48 | .233 | .038 |

| Effect | df | F | *p* | $\eta^2p$ (Effect Size) |
| --- | --- | --- | --- | --- |
| Time (Immediate vs. Delayed) | 1, 79 | 124.09 | <.001 | .620 |
| Time × group | 2,79 | .94 | .39 | .024 |
| Type image × Time | 1, 79 | 1.93 | .169 | .025 |
| Type image × time × group | 2,79 | 1.12 | .33 | .025 |

Note. The table shows the mixed effects ANOVA SPSS results of the behavioural memory results (hits minus false alarms of emotional images vs neutral images in time point one or immediate vs time point two or delayed). df = degrees of freedom, F = F-statistic (from SPSS output), *p* = p-value (significance level), $\eta^2p$ (Partial Eta Squared) = Effect size (SPSS reports this in "Tests of Within-Subjects Effects").

A 2 (type of image memorized: Neutral, Emotional) × 2 (time: Immediate, Delayed) × 3 (Group: Young, Old, MCI) mixed effects ANOVA revealed a significant main effect of type of image memorized, F(1, 79) = 5.46, p = .022, indicating that emotional images were better remembered than neutral ones. There was also a significant main effect of time, F(1, 79) = 124.09, p < .001, suggesting that immediate recall was superior to delayed recall. However, the type of image memorized × time interaction was not significant, F(1, 79) = 1.93, p = .16. A main effect of group was found, F(2, 79) = 30.66, p < .001, with post hoc Tukey tests indicating that MCI participants had significantly lower memory scores compared to both YA (mean difference: .40, p < .001) and OA (mean difference: .31, p < .001) groups, while YAs and OAs did not differ (mean difference: .09, p =.206). Interactions are reported in table 3.

Regarding the delayed recognition of neutral images we found a positive correlation in the whole group between LC integrity and recognition of neutral images (n = 82, r = .373, p <.001). We only found a positive correlation between these variables in the MCI group (n = 24, r = .408, p = .048). When looking at the emotional images, we only found a positive correlation between LC integrity and delayed recognition memory of emotional images when looking at the whole group (n = 82, r = .357, p <.001).

### 3.2. Structural LC results: integrity

As shown in Figure 3, LC integrity measured as the contrast ratio between the LC and a region of the pons differed between all three participant groups (F(2, 79) = 14.38, p < .001) and was larger in older adults compared to younger adults (mean difference = 0.03, p = .02), larger in older adults as compared to age-matched individuals with MCI (mean difference = 0.06, p < .001), and larger in younger adults as compared to individuals with MCI (mean difference = 0.03, p =.02).

**FIGURE 3.**
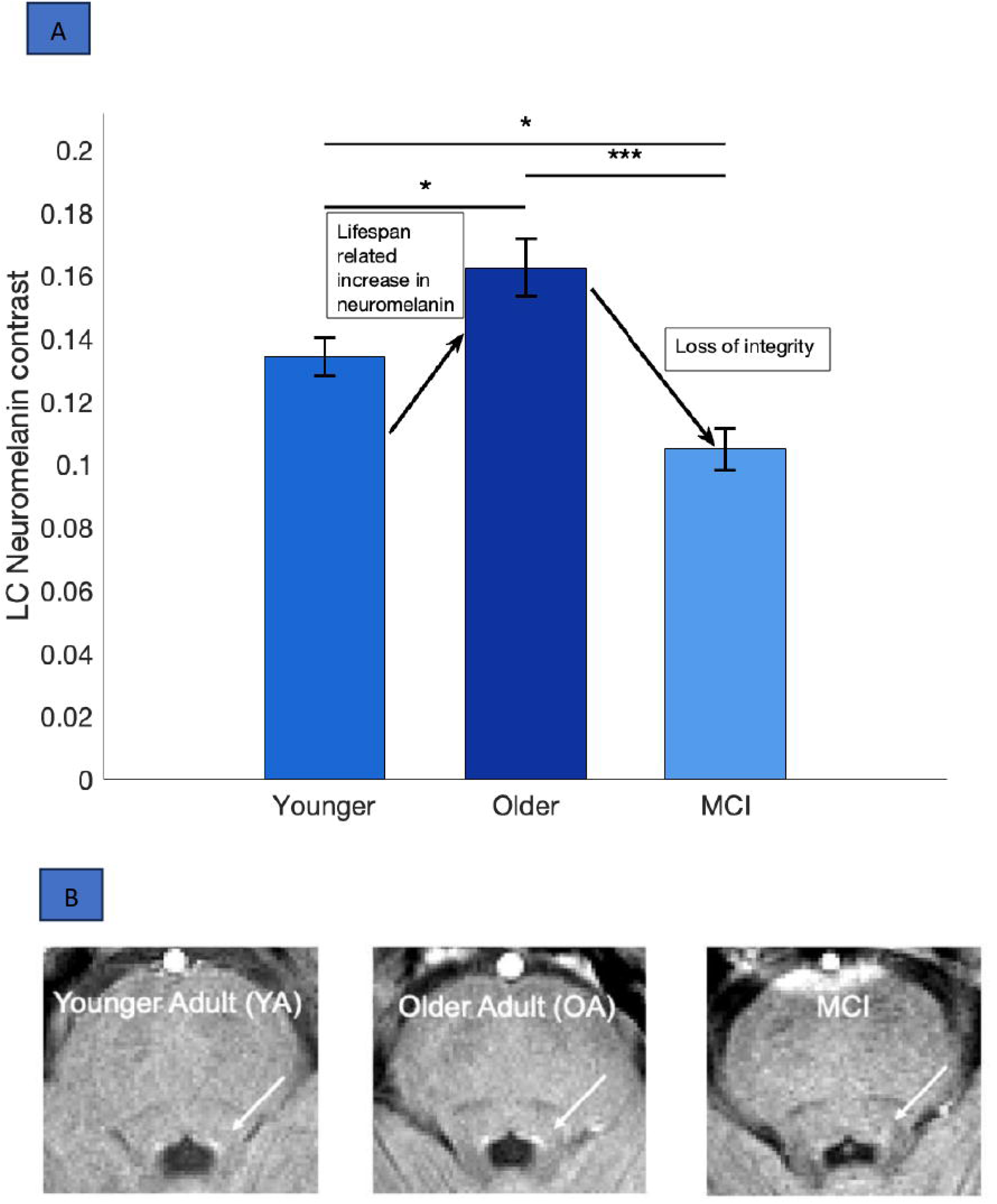
**A.** Boxplots of the statistical difference in LC integrity across groups. Lines with asterisks indicate statistically significant differences (younger adults < older adults; younger adults > MCI and older adults > MCI). **B.** Example structural LC scans for one younger adult, one older adult and one MCI patient, shown in the mean across 3 repeated neuromelanin-sensitive scans. The LC is apparent as hyperintense area (white arrow).

### 3.3. Task fMRI results

#### 3.3.1. Sanity check

Increased BOLD signal in motor areas were explored as a ‘sanity check’ to assess whether activated areas in the three groups were affected by examining activations in a joint anatomical space across YAs, OAs, and MCI patients. They were assessed through a second-level analysis of the contrast ’response larger than fixation cross,’ to examine brain BOLD activations that are stronger when participants press any mouse button compared to when they passively view a baseline fixation cross. Note that these activations will also contain visual effects as responses were given during scene stimuli which are more complex visually than fixation crosses.

On the study template, in the whole group (i.e. in a joint anatomical space across all three groups (YAs, OAs, MCI together), as expected, the contrast ’response larger than fixation cross’ revealed greater activation in motor regions such as the supplementary motor cortex, precentral gyrus, as well as the parahippocampus, likely reflecting scene processing related effects. There were some differences across groups, with the precentral gyrus and supplementary motor cortex activated only in the YA group, while the parahippocampus showed activation across all three groups.

Importantly, the activated ROIs of the ‘response larger than fixation cross’ contrast remained largely consistent when examined in the group-specific templates, except for the parahippocampus in the YA group, which did not show a significant activation (see Supplementary Material Table 1 and Supplementary Material Figure 2.1). This suggests that activations for the three participating groups can be examined and compared in the whole - group anatomical space.

#### 3.3.2. Negative vs neutral images

The one-sample T-test results (study-specific template; Supplementary Material Table 2 and Figure 4 in this main manuscript) in the whole group analysis showed increased activation in regions associated with emotional processing, including caudate, amygdala left, mid / posterior cingulate cortex, the thalamus and the inferior frontal cortex. Unexpectedly, the lingual gyrus and the primary visual cortex also showed heightened activation.

**FIGURE 4.**
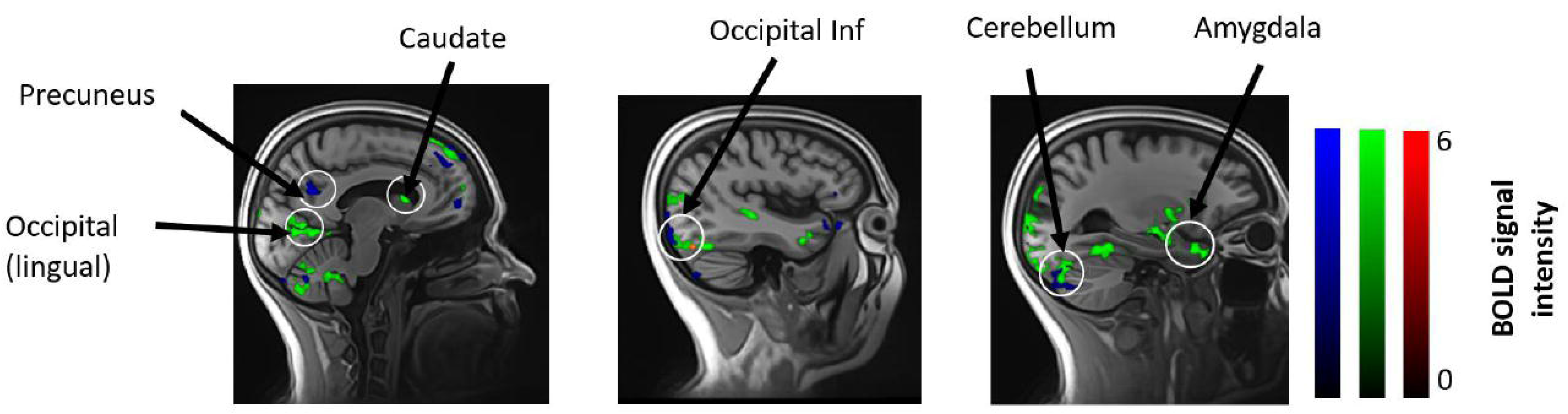
Activations of the contrast Emotional > Neutral scenes in the study template. Activations of the younger group (e.g., amygdala, cerebellum, caudate) in green, of the older adult group (e.g., occipital and temporal regions) in blue, and of the MCI patients (occipital inferior) in red.

When assessing the groups separately, the YA group showed increased activation in the insula, caudate, amygdala, hippocampus and the occipital inferior cortex. Occipital and temporal regions were activated in each group. Between-group comparisons revealed no significant differences in activation between YA vs. OA and OA vs. MCI (an overview of the results is displayed in Table 4 of this manuscript as well as in Supplementary Materials Table 2, and in Figure 4 in the main manuscript).

**Table 4.**
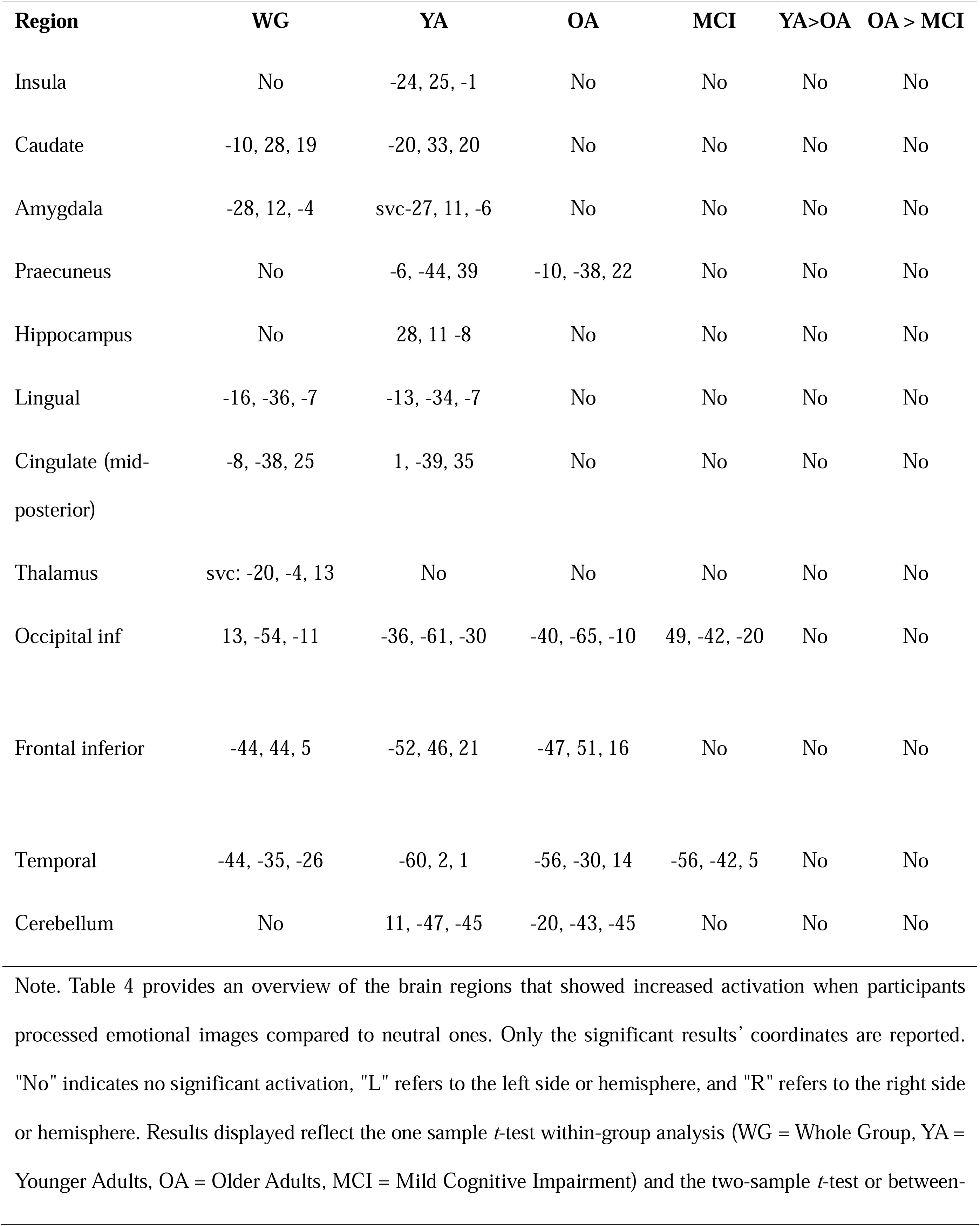
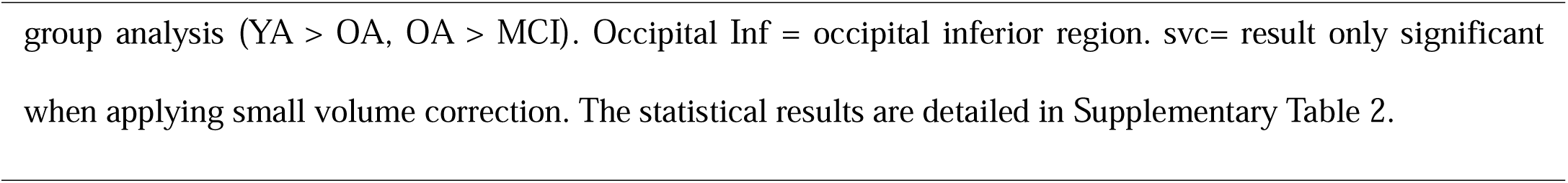
Emotional processing larger than neutral using the study template.

| **Region** | **WG** | **YA** | **OA** | **MCI** | **YA>OA** | **OA > MCI** |
| --- | --- | --- | --- | --- | --- | --- |
| Insula | No | -24, 25, -1 | No | No | No | No |
| Caudate | -10, 28, 19 | -20, 33, 20 | No | No | No | No |
| Amygdala | -28, 12, -4 | svc-27, 11, -6 | No | No | No | No |
| Praecuneus | No | -6, -44, 39 | -10, -38, 22 | No | No | No |
| Hippocampus | No | 28, 11 -8 | No | No | No | No |
| Lingual | -16, -36, -7 | -13, -34, -7 | No | No | No | No |
| Cingulate (mid-posterior) | -8, -38, 25 | 1, -39, 35 | No | No | No | No |
| Thalamus | svc: -20, -4, 13 | No | No | No | No | No |
| Occipital inf | 13, -54, -11 | -36, -61, -30 | -40, -65, -10 | 49, -42, -20 | No | No |
| Frontal inferior | -44, 44, 5 | -52, 46, 21 | -47, 51, 16 | No | No | No |
| Temporal | -44, -35, -26 | -60, 2, 1 | -56, -30, 14 | -56, -42, 5 | No | No |
| Cerebellum | No | 11, -47, -45 | -20, -43, -45 | No | No | No |

Note. Table 4 provides an overview of the brain regions that showed increased activation when participants processed emotional images compared to neutral ones. Only the significant results' coordinates are reported. "No" indicates no significant activation, "L" refers to the left side or hemisphere, and "R" refers to the right side or hemisphere. Results displayed reflect the one sample *t*-test within-group analysis (WG = Whole Group, YA = Younger Adults, OA = Older Adults, MCI = Mild Cognitive Impairment) and the two-sample *t*-test or between-

After controlling for the covariates age, grey matter volume (GMV), and LC integrity, some results in the emotional > neutral contrast varied. In the YA group, when controlling for age and GMV, the results were no longer significant. Conversely, most results remained intact after controlling for LC integrity, except in the lingual gyrus, insula, and fusiform. In the OA group, no significant activations were observed when controlling for age and GMV. However, when controlling for LC integrity, all results remained stable except in the middle temporal, inferior temporal, and precuneus regions. Finally, the two regions that showed increased activation in the emotional contrast in MCI (i.e., inferior temporal and inferior occipital) did not show significant activations after controlling for any of the covariates.

Taken together, group-wise analyses revealed that only the YAs showed stronger activations in the insula, caudate, cingulate, amygdala and the hippocampus for emotional as compared to neutral stimuli, while all groups showed activations in the occipital inferior, suggesting enhanced visual processing of emotional stimuli and temporal regions. No significant between-group differences were found when comparing YAs vs. OAs, YA vs MCI or OAs vs. MCIs. These results suggest that while the networks engaged by emotionally salient stimuli are similar across age groups, the strength of activation does not differ significanly. Age and atrophy seem to be explaining a lack of specific effect for emotional processing (see Table 4).

#### 3.3.3. Remembered images larger than not remembered images

In the whole group, the contrast “remembered images larger than not remembered images”, revealed increased activation only in the right putamen ( p_FDRc- cluster corr) *p*=.05). When assessing the groups separately the YA showed an increased activation in the parahippocampus right and left (p_FDRc-cluster corr) (*p* <.001), as well as in the precentral gyrus, cuneus and precuneus. The OA showed increased activation in the right and left putamen (*p* = 0.04 and *p*= .05 respectively) and a trend in the left LC small volume corrected (x = -6, y = -8, z = -27; p_FWEc=._08, k_E_= 4). No significant activations were observed in the MCI group. The lack of significant encoding-related activation in MCI is consistent with their behavioural memory impairment and suggests a failure to engage the regions necessary for successful encoding.

When comparing results between pairs of groups, the precentral, cerebellum area 7b and precuneus showed an increased activation in the YA in contrast to the OA. The LC was slightly more active in the OA as compared to the MCI group. The cuneus, fusiform and precentral showed an increased activation in YA > MCI in successful remembered > not successfully remembered.

There were no significant differences in the contrast “remembered larger than not remembered” in the following group comparisons: OA larger than YA, MCI larger than OA and MCI larger than YA. An overview of the results is shown in Table 5 and Figure 5 of this main manuscript as well as in Supplementary Table 3.

**FIGURE 5.**
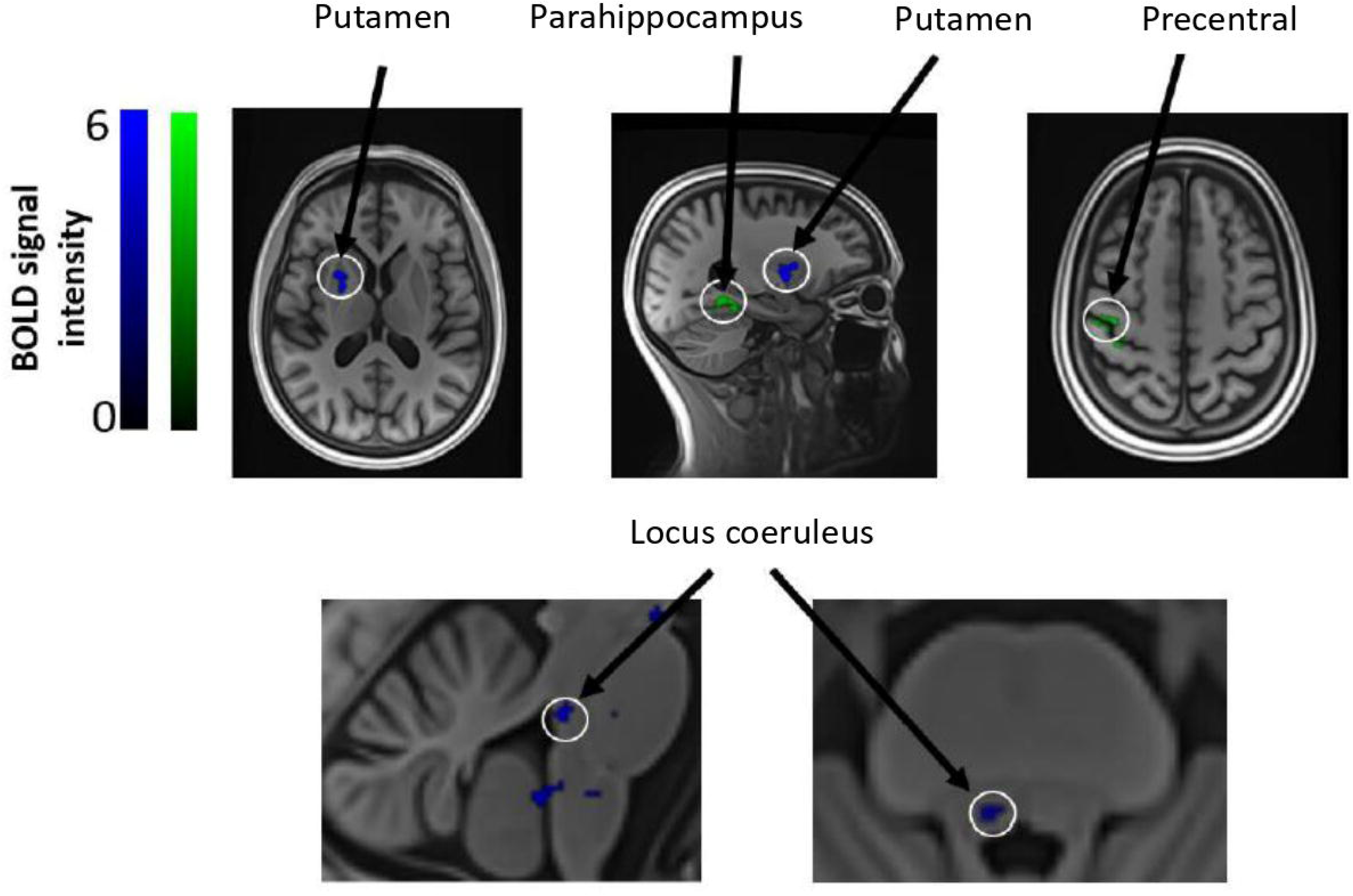
Activations of the contrast remembered > not remembered in the study templates. Activations of the younger group (e.g. parahippocampus) in green, of the older adult group (e.g. putamen) in blue. No significant activations were observed in MCI patients.

**Table 5.** Remembered images larger than not remembered images using the study template.

| **Region** | **WG** | **YA** | **OA** | **MCI** | **YA>OA** | **OA > MCI** |
| --- | --- | --- | --- | --- | --- | --- |
| Putamen | svc:19, 29, 6 | Svc:18, 25, 5 | -23, 32, 8 | No | No | No |
| Caudate | No | No | No | No | No | No |
| Parahippocampus | svc-32, 1, -17 | 36, -13, -9 | No | No | No | No |
| Precentral | No | 35, -6, 41 | No | No | 35, -1, 54 | No |
| Precuneus | No | No | No | No | 11, -58, 34 | No |
| Occipital mid | No | -21, -72, -5 | No | No | No | No |
| Lingual | No | -29, -25, -16 | No | No | No | No |
| LC | No | No | svc-6, -8, -27* | No | No | svc-6, -8, -27* |

Note. Table 5 provides an overview of the brain regions that showed increased activation of the images displayed during the scan session that were subsequently remembered in the memory recognition test.
Coordinates of the significant results are provided. "No" indicates no significant activation, "L" refers to the left side or hemisphere, and "R" refers to the right side or hemisphere. Results displayed reflect the one sample *t*-test within-group analysis (WG = Whole Group, YA = Younger Adults, OA = Older Adults, MCI = Mild Cognitive Impairment) and the two-sample *t*-test or between-group analysis (YA > OA, OA > MCI). Caudate: increased in OA only after controlling for covariates. \*LC: Locus Coeruleus in OA and OA > MCI only marginal significant and not significant after controlling for covariates. Svc = results significant only when small volume corrected. The specific statistical results are detailed in supplementary table 3.

Most of the results of the contrast “remembered larger than not remembered” remained significant after controlling for the covariates “LC integrity”, “GMV” or “age” with some exceptions: Within the older adult (OA) group, controlling for “age” and “GMV” revealed significant activation in the caudate (*p* = .04), which was not observed in the initial analysis. This suggests that accounting for interindividual differences in age and atrophy captures variance which occluded identifying larger activations in the caudate for remembered as compared to not remembered stimuli.

Conversely, in the OA > MCI comparison, the LC was no longer significantly activated in the remembered > not remembered contrast after controlling for age and LC integrity. This suggests that interindividual differences in age and LC integrity contribute to differences in LC function during memory encoding.

In conclusion, our results suggest that parahippocampus, precentral gyrus and cuneus are implicated in successful memory encoding in the YA and differentiate memory encoding from the older group (OA group and MCI), while the putamen, the caudate and the LC are key in healthy aging.

## 4. DISCUSSION

The main objective of this study was to explore aging and MCI changes in immediate and delayed episodic memory performance, associated brain functional activations related to the processing of neutral and emotional scenes and the impact of LC integrity in our outcomes.

Behavioral results showed comparable memory performance between younger and older adults, likely due to the comparatively easy memory task necessary to allow a comparative assessment with the MCI patient group. As expected, the MCI group showed poorer memory. Moreover, immediate recognition outperformed delayed, and emotional images were better remembered than neutral ones. However, no group by emotional vs. neutral memory effect was observed.

The amygdala, a region that has been consistently linked with processing emotionally arousing stimuli [48–50] showed an increased activation only in the whole group data and in the YA during the processing of negative emotional as compared to neutral images. Despite no significantly larger activations in amygdala in OAs or MCI patients, during emotional as compared to neutral scenes, no significant age effects were observed in the preference for emotional as compared to neutral images[51].

Several studies have investigated the relationship between subjective arousal ratings and objective physiological responses in OA, revealing discrepancies. For instance, one study [52] found that OA reported higher subjective arousal to negative images compared to YA, yet their electrodermal activity was significantly lower. Similarly, [53] observed that OA rated negative images as more arousing than YA did, but their physiological responses did not correspondingly increase. This suggests a disconnect between subjective experience and objective arousal in OA, highlighting the need to include both self-report and physiological data (e.g., electrodermal activity, pupil dilation) in future work.

Apart from the relevance of the amygdala previously reported as a key region in emotional processing and memory encoding, further brain regions are assumed to additionally contribute. In line with prior work [54,55], the present study showed that the medial prefrontal cortex, inferior frontal cortex, inferior temporal cortex, striatal temporal cortex, lateral occipital cortex and medial dorsal nucleus of the thalamus are activated in emotional scene processing. The insula, also typically associated with emotional processing [56], showed an increased activation in the YA. We also observed enhanced activation in the lingual gyrus (YA), precuneus, and primary visual cortex (all groups except MCI). Although not classic emotion regions, previous studies found higher visual cortex activation with emotional images, possibly reflecting greater attentional focus [54,57].

In the contrast ’remembered larger than not remembered,’ the putamen was consistently activated in all three groups. As expected, the parahippocampus showed increased activation, albeit only in YA; this effect was absent in OA and MCI, likely due to greater atrophy and loss in function in older individuals.

Our results suggest, furthermore, that the precentral sulcus and the cuneus are involved in successful memory encoding in young participants. These findings align with previous studies that found that motor regions are activated during memory tasks that require a response action such as pressing a button [58]. A meta-analysis further identified that the precentral sulcus as part of the working memory network [59] and both the precentral sulcus and cuneus have been seen to be involved in visual and working memory[59,60]. A lack of activation in these regions is shown in the OA and MCI groups, suggesting an ageing-related decline in their contribution. Conversely, the fusiform, which has been linked with recognition of objects and faces [55,61,62], differentiates YA from MCI, as the former group shows a higher activation.

Given the known role of the cuneus in visual processing and visual encoding [59,63], its activation is likely related to the processing of the scene stimuli. This process is crucial to forming connections with other brain memory related regions like the hippocampus, the fusiform gyrus and the amygdala, thereby strengthening memories. Reduced activation in these areas in OA and MCI may contribute to their poorer memory performance. In addition to the cuneus, the precuneus, a region often associated with episodic memory, showed increased activation in successful memory encoding in YA in contrast to the OA. Aligned with these results, a previous study indicated that highly vivid memories were associated with an increased activation in the precuneus in YA in contrast to OA[64]. Additionally, a review of fMRI and PET studies exploring the functional and behavioral role of the precuneus concluded that increased activation of this region is associated with higher memory performance [65]. Interestingly, these findings were found not only in studies using a classic lab set of object stimuli, but also other type of naturalistic stimuli that enable to assess episodic and autobiographic memory, e.g., family images. This implies that the precuneus is activated when encoding any of these type of lab stimuli (scenes and objects) and it is associated with episodic memory, impaired in early stages of amnestic MCI.

One of our main goals was to shed light on the functional role of the LC in memory encoding.The LC showed a slightly increased activation (trend) in the OA as compared to MCI, and did not differ in activation from YA. A recent study showed that the LC functional activation in OA in contrast to YA seems to be increased, especially when exposed to negative emotional images [25]. This increased activation may reflect compensation for age-related structural LC decline, which might no longer be possible in MCI patients with more advanced structural LC decline as in our study. This hypothesis should be further investigated with other samples.

Finally, results that were significant or showed a trend at a threshold of *p* < .005 at the voxel level and *p* < .001 at the cluster level did not remain significant after controlling for GMV and LC integrity, suggesting that interindividual differences in LC function are closely tied to interindividual differences in its structure. LC integrity and GMV appear to influence memory performance and LC activation during incidental encoding, regardless of diagnosis.

MCI showed no significant activations, matching low recognition memory and limited encoding. The overall poor performance in MCI could be related to lower LC integrity.

Taken together, LC integrity was associated with lower brain activity during encoding and emotional salience processing but the relationship with delayed memory performance in MCI was not clear.

Future studies should include patients in different stages of MCI (i.e., with different degrees of cognitive impairment and AD pathology) and add CSF and blood-TAU / betamiloid parameteres , to explore whether our findings remain consistent.

In conclusion, our study suggests that:

- The beneficial effect of emotional salience on memory is preserved across all groups despite the overall memory decline in MCI. This preserved effect in the behavioural data is consistent with the finding that all groups showed heightened activation in occipital and temporal regions during emotional processing.
- YA shows stronger activation in amygdala and insula when viewing negative emotional images while OA and MCI recruited temporal and occipital regions. These activations are not significant when comparing across groups.
- Successful encoding involve increased activation of the parahippocampus, precentral gyrus, and precuneus in YA; and caudate, putamen, and an LC trend in OA and in OA larger than in MCI.
- The MCI group shows no significant activation during successful encoding. A lack of LC activation in LC seems to be related to overall poorer performance but not to emotional valence. Observations in the literature of hyperactivity in MCI as a compensatory mechanism due to TAU accumulation and atrophy in some encoding related structures cannot be explained by LC activity.
- LC integrity is reduced in MCI compared to OA, and a trend for higher LC activation in OA than MCI suggests a potential role of the LC in supporting memory encoding in healthy ageing. These results were not significant when controlling for age and LC integrity. Meaning that the activation of the LC is dependent of LC integrity but independent of an MCI diagnosis. Further studies should include MCI patients in an early and late stage and add CSF and blood-TAU / betamiloid parameteres, to explore whether our findings remain consistent.

Limitations:

This study has the following limitations. First, the modest sample size may affect the generalizability of results, especially between MCI and healthy aging groups. Second, participant characterization lacked biological markers (e.g., CSF tau, amyloid, genetic risk), limiting insights into underlying pathology. Third, we did not assess subjective arousal or include physiological measures like electrodermal activity or eye-tracking, which could clarify attentional and emotional engagement. Lastly, image memorability factors such as distinctiveness or familiarity were not evaluated, which may also influence encoding success in addition to emotional valence.

## ACKNOWLEDGEMENTS

The authors would like to give a special thanks to all the participants who kindly provided their time and effort.

## SOURCES OF FUNDING

This work was supported by:

Wellcome Trust. Gibbs Building 215 Euston Road, London NW1 2BE. Grant ref.: 203147/Z/16/Z;

The Department of Psychology, University of Innsbruck, A-6020, Innsbruck, Austria;

Institute of Cognitive Neurology and Dementia Research, Otto-von-Guericke-University Magdeburg, D-39120, Magdeburg, Germany;

Institute of Cognitive Neuroscience, University College London, London, UK-WC1E 6BT, UK;

CBBS Center for Behavioral Brain Sciences, D-39120, Magdeburg, Germany; and, German Center for Neurodegenerative Diseases (DZNE), D-39120, Magdeburg, Germany.

## CONSENT STATEMENT

Informed consent was obtained from all subjects involved in this study.

## DATA AVAILABILITY STATEMENT

Further details on the data are available upon request. Please contact Lucía Penalba-Sánchez at the Institute of Cognitive Neurology and Dementia Research (IKND), German Center for Neurodegenerative Diseases (DZNE), Magdeburg.

## Abbreviations

AD: Alzheimeŕs Disease
MCI: amnestic Mild Cognitive Impairment
OA: Older Adults
YA: Younger Adults
LC: Locus Coeruleus
GMV: Gray Matter Volume.

