## Supplementary material for "Differences in LC integrity and fMRI activations in healthy aging and MCI associated with successful memory encoding"

**Figures:**

**Figure 1.** Histograms of in-plane distances between landmarks defined on the study-specific template and single-subject landmarks delineated on mean functional images transformed into the template space. Each inset in the corresponding histogram plot indicates its anatomical position on the structural MNI template. The detailed procedure for selecting and placing the landmarks, as well as quantifying the distances, is described in Yi et al.'s (2023) work. Note that the distances in the Outline Brainstem landmarks vary, as they were placed anywhere along the outline of the brainstem border. The mean and standard deviation distances for landmarks are described underneath the anatomical position insets.

#### 1.1. Histograms landmarks (study template)

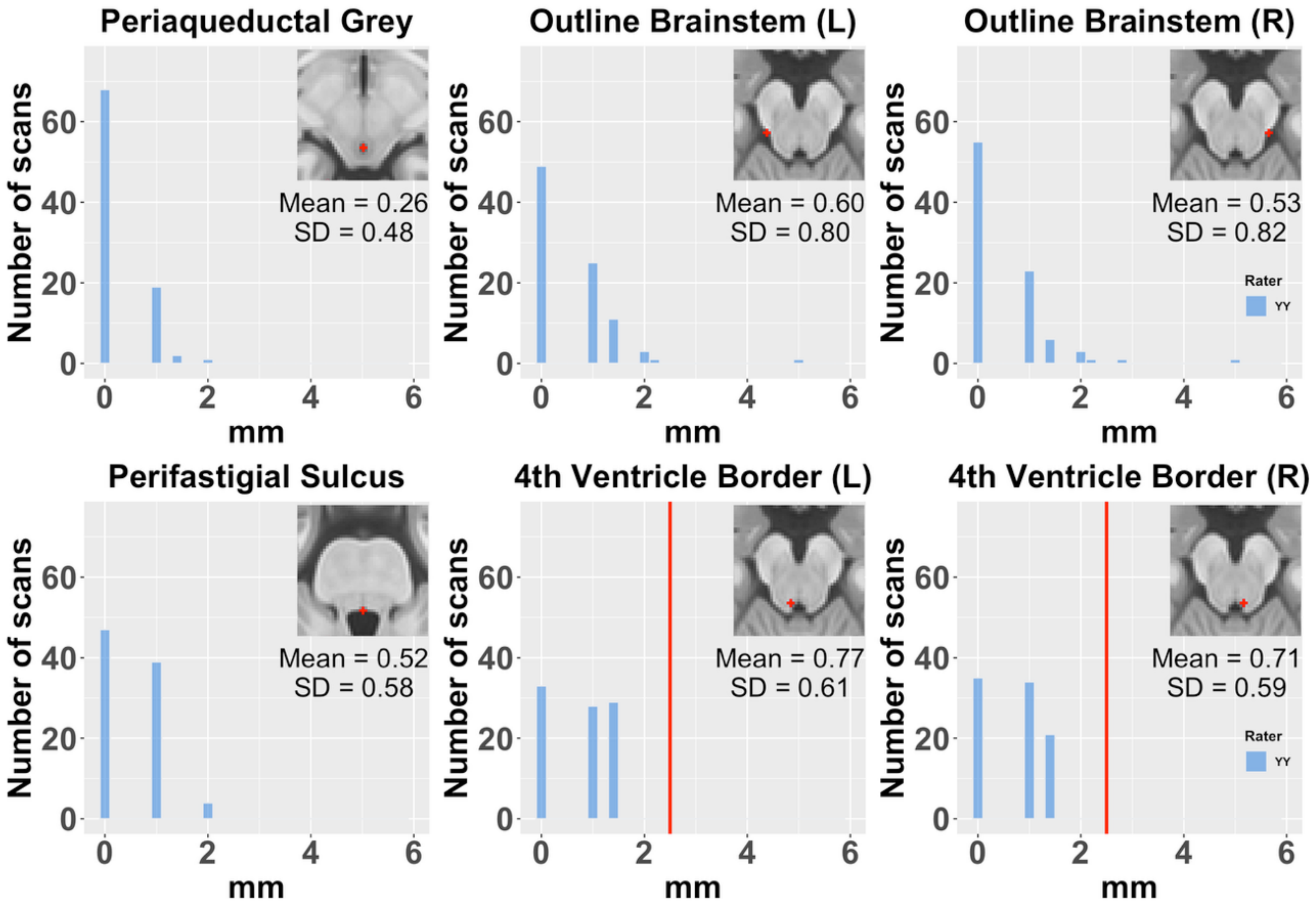


2.

**1.2. . Histograms landmarks (younger adults' template)**

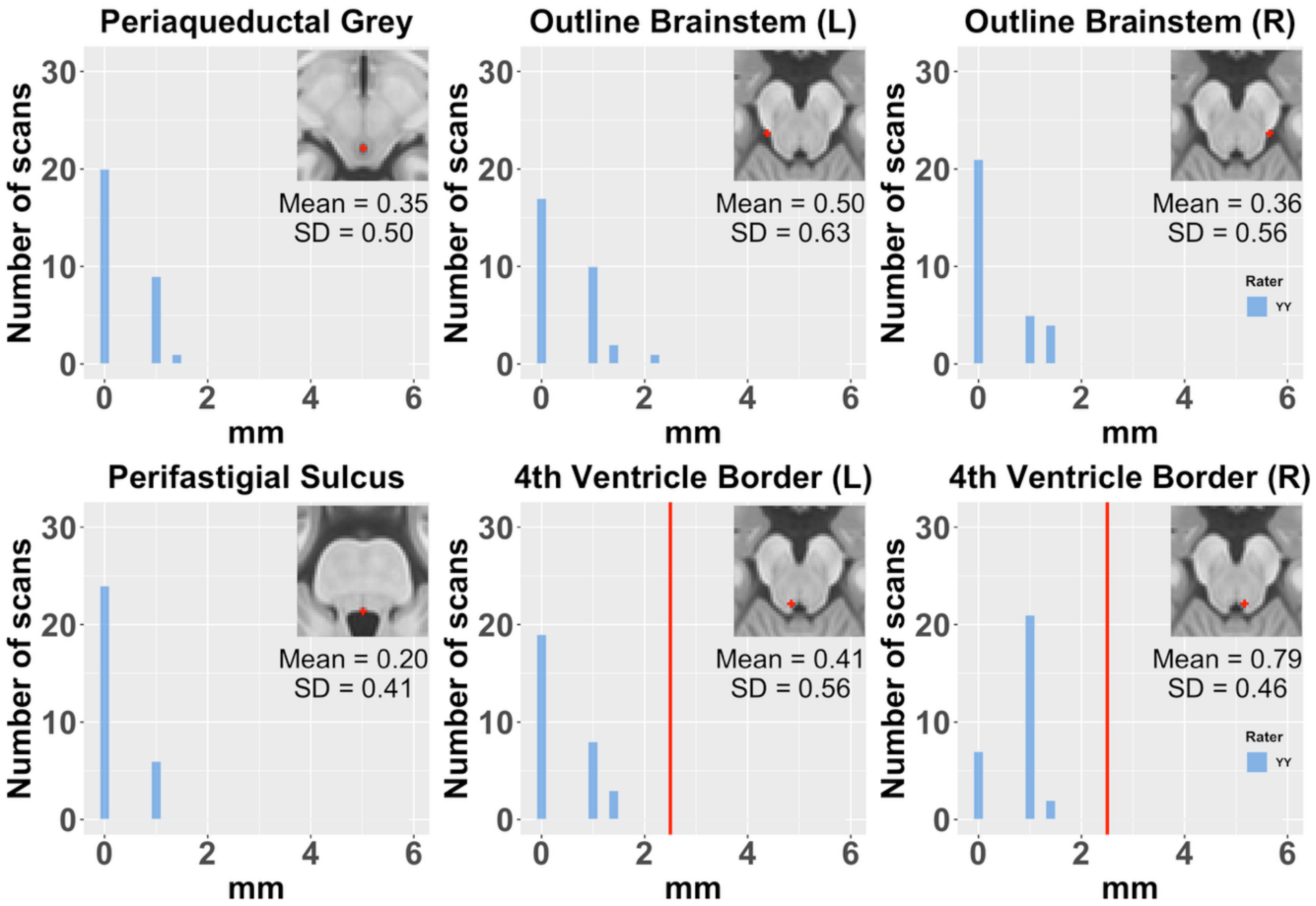

**1.3. . Histograms landmarks (older adults' template)**

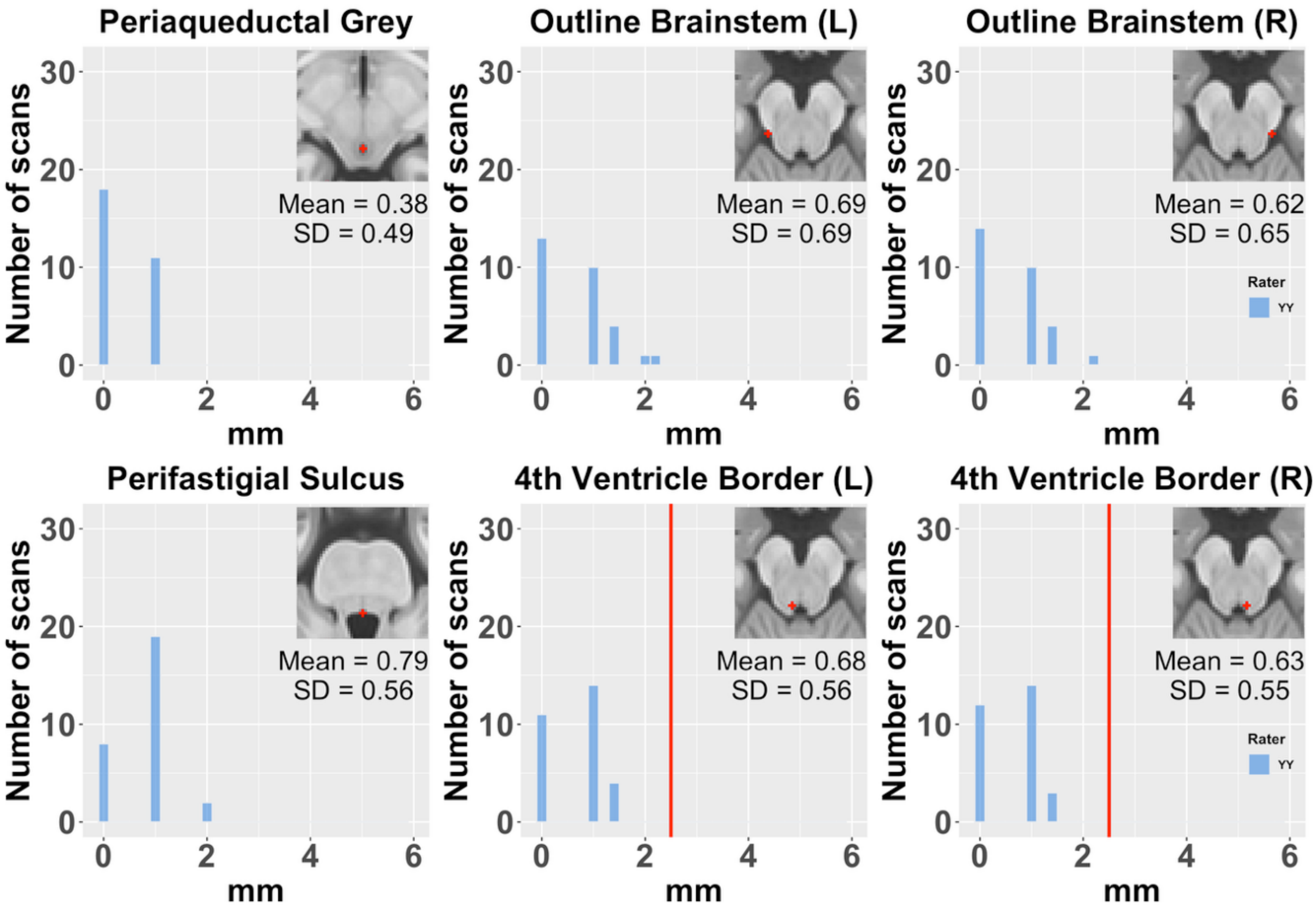

#### 1.4. Histograms landmarks (MCI template)

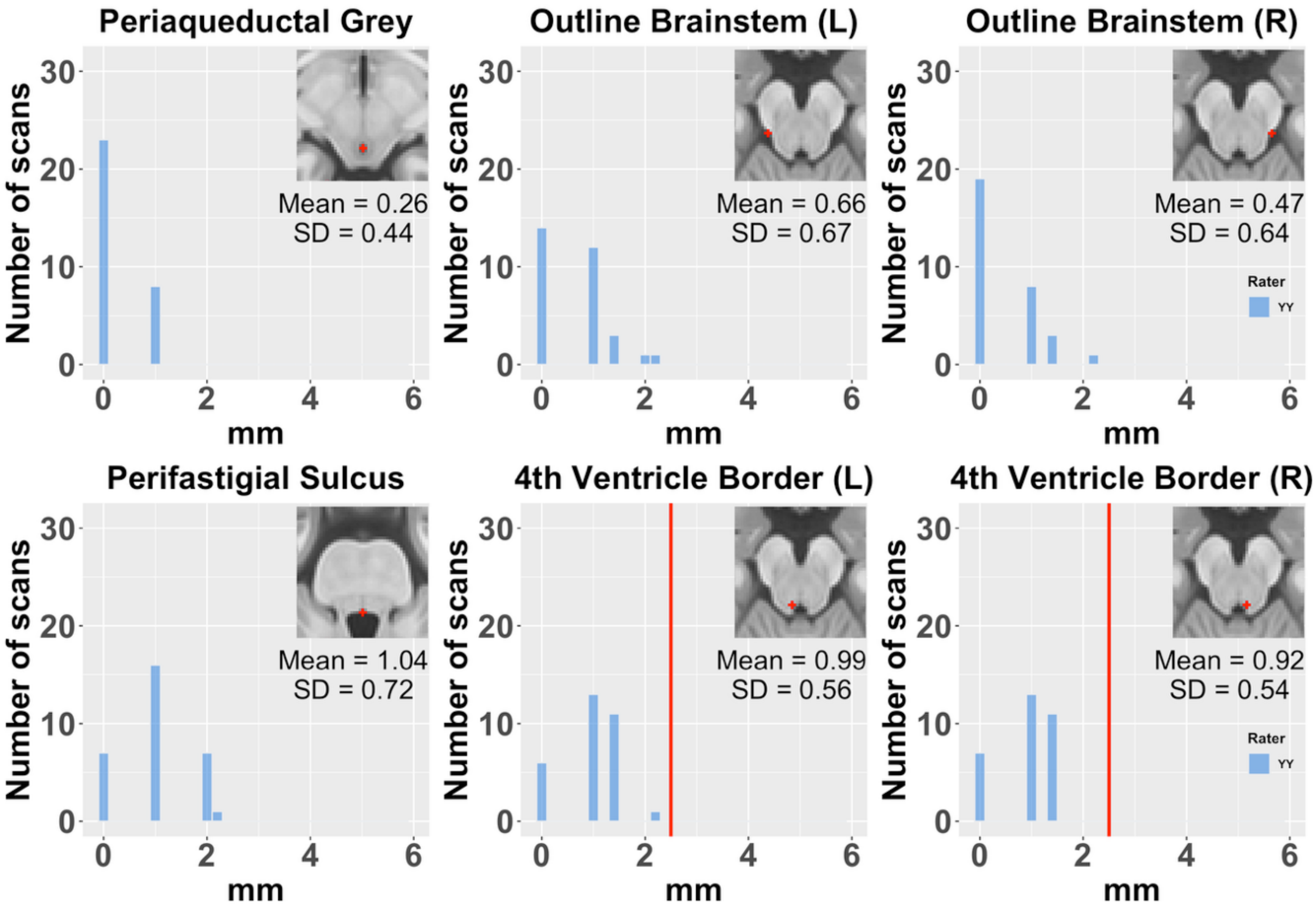

**Figure 2. Sanity check**

Functional brain activations of the sanity check motor contrast (response > fix). Results are compared in study template across all three groups (top left), in study template space for individual group results (top right) as well as in group-specific templates (bottom). Activations in younger adults are indicated in green, in older adults in blue, and in MCI in red. Sanity checks examined differences in activated brain areas in group-specific and study template space which would suggest distortions due to merging across group-specific anatomies. Group-specific activation patterns were comparable in study and group-specific space, suggesting that between group comparisons in study space are not strongly affected by anatomical differences between groups (Supplementary table 1).

**2.1.Response > fixation cross** (study template)

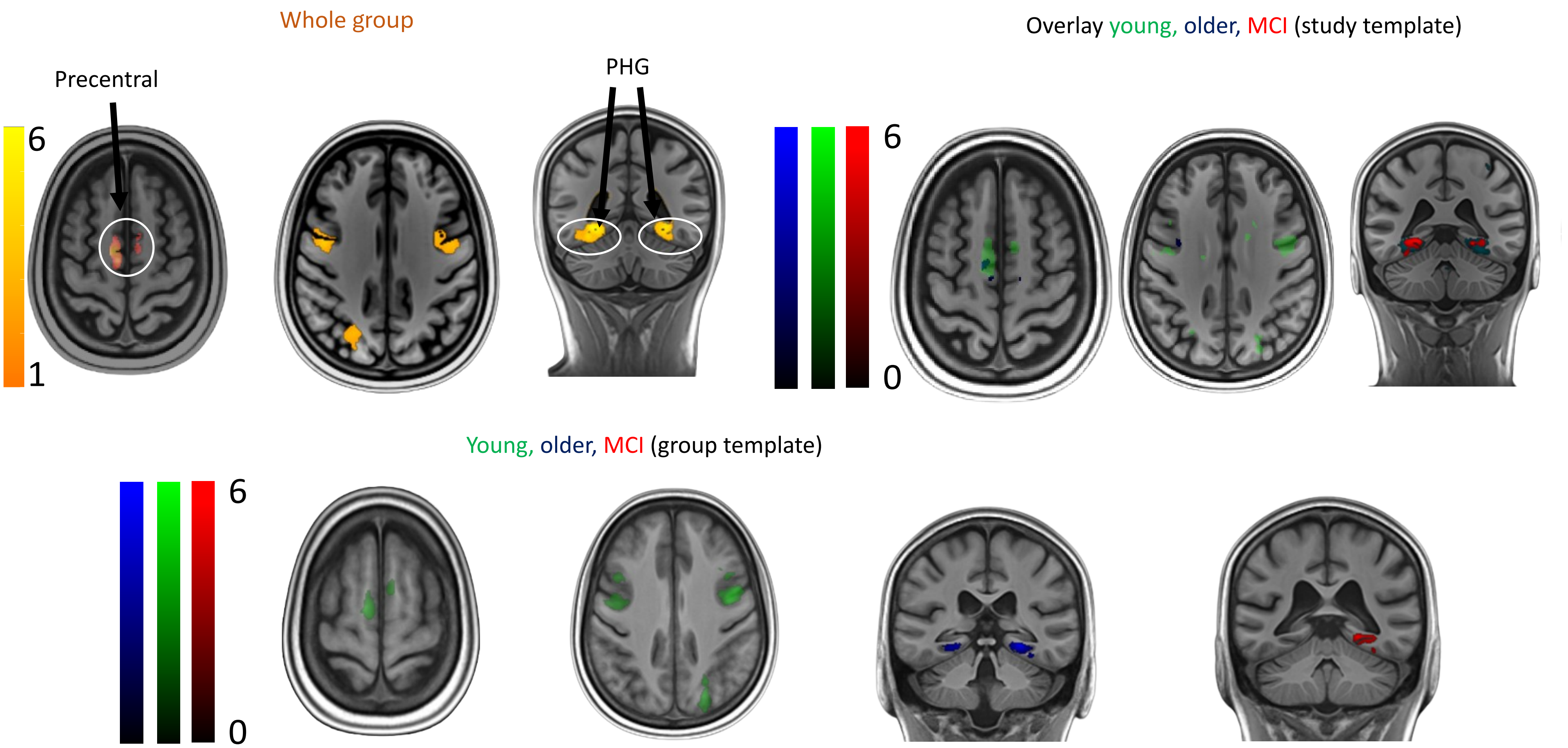

### Discussion sanity checks

As expected, motor-related brain regions exhibited enhanced event-related BOLD activation in the sanity check. For instance, the supplementary motor cortex and precentral gyrus were activated during finger-tapping tasks, particularly in the younger group. Interestingly, the parahippocampus was additionally significantly activated in all three groups, likely reflecting its role in scene processing. As the majority of areas identified in group-specific templates are also observed in the study template, it can be assumed that anatomical distortions in bringing individuals in the joint study template space is not affecting the interpretability of between group comparisons in study space substantially.

### Tables:

**Functional activations contrasts of interest. Uncorrected p values, threshold p<.005, cluster p(FDR) corrected. When applying small volume corrections, threshold p <.01 p(FEW-corrected)**

**Table 1. Sanity check response larger than fixation cross (motor)**

| **Motor whole group study template (cl 491)** | | | | | | |
|---|---|---|---|---|---|---|
| **Region** | **p(FWE-corr)** | **p(FDR-corr)** | **equivk** | **x** | **y** | **z** |
| Parahippocampus | <.001 | <.001 | 36832 | 30 | -22 | -12 |
| | | | | 22 | -12 | -13 |
| | | | | 32 | -40 | -19 |
| | <.001 | <.001 | 22979 | -30 | -19 | -12 |
| | | | | -1 | 25 | 25 |
| | | | | -29 | -33 | -15 |
| Precuneus | 0.169 | 0.007 | 610 | 20 | -38 | 7 |
| | | | | 25 | -40 | 15 |
| Precentral (motor) | 0.002 | <.001 | 1236 | 42 | 16 | 35 |
| Sup motor area | 0.001 | <.001 | 1420 | -8 | 5 | 57 |
| | | | | -5 | 13 | 52 |
| | 0.404 | 0.016 | 491 | 9 | 12 | 61 |
| | | | | 8 | 16 | 54 |
| | | | | -9 | -3 | 63 |

| | | | | | | |
|---|---|---|---|---|---|---|
| Pineal body | 0.256 | 0.01 | 555 | 4 | -8 | -3 |
| | | | | -1 | 1 | 2 |
| | 0.015 | 0.001 | 932 | 33 | 20 | -23 |
| | | | | 38 | 17 | -17 |
| Precentral | 0.016 | 0.001 | 927 | -42 | 14 | 35 |
| | | | | -34 | 21 | 35 |
| | 0.004 | <.001 | 1143 | -37 | -6 | 49 |
| | | | | -36 | 1 | 44 |
| | 0.142 | 0.006 | 633 | 42 | -1 | 48 |
| | | | | 36 | 5 | 46 |

**Motor younger adults study template (cl 332)**

| **Region** | **p(FWE-corr)** | **p(FDR-corr)** | **equivk** | **x** | **y** | **z** |
|---|---|---|---|---|---|---|
| Parahippocampus | <.001 | <.001 | 28773 | 29 | -24 | -13 |
| | | <.001 | | 32 | -40 | -19 |
| | | <.001 | | 27 | -47 | -21 |
| | <.001 | <.001 | 20056 | -29 | -31 | -17 |
| | | <.001 | | -27 | -47 | -19 |
| | | <.001 | | -31 | -19 | -12 |
| Precentral | <.001 | <.001 | 1534 | 43 | 20 | 31 |

| | | | | | | |
|---|---|---|---|---|---|---|
| | <.001 | <.001 | 1567 | 43 | 1 | 47 |
| | | | | 35 | 5 | 46 |
| | | | | 36 | -7 | 49 |
| | 0.001 | <.001 | 1198 | -40 | 13 | 36 |
| | | | | -51 | 12 | 33 |
| | | | | -37 | 21 | 35 |
| | <.001 | <.001 | 1855 | -42 | -5 | 47 |
| | | | | -34 | -10 | 49 |
| | | | | -32 | 2 | 54 |
| Superior motor area | <.001 | <.001 | 1690 | -6 | 18 | 52 |
| | | | | -7 | 5 | 58 |
| | | | | -12 | -1 | 61 |
| Superior motor | 0.251 | 0.008 | 502 | 9 | 16 | 54 |
| Frontal mid | 0.019 | 0.001 | 812 | 40 | 37 | 28 |
| | 0.354 | 0.011 | 459 | -40 | 36 | 31 |
| | | | | -37 | 39 | 24 |
| | | | | -37 | 26 | 31 |
| Fusiform / lingual | 0.647 | 0.023 | 373 | 35 | 3 | -18 |
| | 0.374 | 0.011 | 452 | 35 | 20 | -21 |
| | | | | 28 | 27 | -22 |
| | | | | 38 | 11 | -14 |

| | | | | | | |
|---|---|---|---|---|---|---|
| Parietal sup | 0.689 | 0.024 | 362 | -22 | -49 | 38 |
| | | | | -25 | -42 | 32 |

**Motor younger adults group template (cl 300)**

| Region | p(FWE-corr) | p(FDR-corr) | equivk | x | y | z |
|---|---|---|---|---|---|---|
| Lingual V1 | <0.001 | <0.001 | 30485 | 29 | -25 | -7 |
| | | | | 29 | -36 | -8 |
| | | | | 32 | -54 | -3 |
| Lingual | <0.001 | <0.001 | 20890 | -29 | -37 | -10 |
| | | | | -29 | -27 | -7 |
| | | | | -27 | -49 | -11 |
| Precentral | <0.001 | <0.001 | 1599 | 41 | 18 | 33 |
| | <0.001 | <0.001 | 1819 | 42 | 5 | 50 |
| | | | | 29 | -12 | 53 |
| | <0.001 | <0.001 | 2116 | -37 | 5 | 49 |
| | | | | -41 | -3 | 52 |
| | | | | -34 | -9 | 54 |
| Precenral mid | <0.001 | <0.001 | 1386 | -41 | 14 | 38 |
| | | | | -52 | 13 | 34 |

| | | | | | | |
|---|---|---|---|---|---|---|
| Supplem motor | 0.149 | 0.005 | 556 | 9 | 17 | 57 |
| | <0.001 | <0.001 | 1687 | -9 | 17 | 56 |
| | | | | -7 | 5 | 62 |
| Frontal mid | 0.005 | <0.001 | 967 | 40 | 38 | 29 |
| | | | | 39 | 32 | 36 |
| | 0.094 | 0.003 | 609 | -40 | 36 | 30 |
| | | | | -38 | 26 | 32 |
| Fusiform | 0.178 | 0.005 | 535 | 36 | 15 | -17 |
| | | | | 41 | 7 | -11 |

**Motor older adults study template (cl 459)**

| **Region** | **p(FWE-corr)** | **p(FDR-corr)** | **equivk** | **x** | **y** | **z** |
|---|---|---|---|---|---|---|
| Fusiform | <.001 | <.001 | 1610 | -29 | -26 | -12 |
| | | | | -30 | -17 | -12 |
| | | | | -21 | -13 | -12 |
| | | | | 23 | -20 | -12 |
| Parahippocampus | <.001 | <.001 | 10855 | 24 | -11 | -13 |

**Motor older adults group template (cl 477)**

| **Region** | **p(FWE-corr)** | **p(FDR-corr)** | **equivk** | **x** | **y** | **z** |
|---|---|---|---|---|---|---|

| | | | | | | |
|---|---|---|---|---|---|---|
| Fusiform | <0.001 | <0.001 | 1566 | -29 | -27 | -14 |
| | | | | -21 | -14 | -14 |
| | | | | -30 | -18 | -14 |
| Parhippocampus | <0.001 | <0.001 | 11188 | 24 | -12 | -15 |
| | | | | 31 | -22 | -14 |
| Fusiform | | | | 31 | -58 | -11 |

**Motor MCI study template (cl 474)**

| Region | p(FWE-corr) | p(FDR-corr) | equivk | x | y | z |
|---|---|---|---|---|---|---|
| Parahippocampus | <.001 | <.001 | 2418 | 25 | -19 | -10 |
| | | | | 24 | -4 | -17 |
| | | | | 21 | -13 | -15 |
| | 0.336 | 0.024 | 474 | -23 | -16 | -14 |
| | | | | -30 | -20 | -13 |
| Occipital Inferior | 0.078 | 0.007 | 651 | 45 | -50 | -7 |
| | | | | 48 | -59 | -7 |
| | | | | 36 | -61 | -4 |

**Motor MCI group template (cl 525)**

| Region | p(FWE-corr) | p(FDR-corr) | equivk | x | y | z |
|---|---|---|---|---|---|---|
| Parahippocampus | <0.001 | <0.001 | 2315 | 24 | -18 | -14 |

| | | | | | | |
|---|---|---|---|---|---|---|
| | | | | 24 | -3 | -18 |
| | | | | 21 | -11 | -18 |
| | 0.189 | 0.014 | 550 | -31 | -20 | -16 |
| | | | | -23 | -15 | -17 |
| Occipital inferior | 0.232 | 0.014 | 525 | 44 | -56 | -11 |
| | | | | 35 | -60 | -8 |

**Table 2. Significant activations of the contrast Emotional Processing (Negative vs. Neutral scenes)**

**Emotional whole group study template (cl 534)**

| **Region** | **p(FWE-corr)** | **p(FDR-corr)** | **equivk** | **x** | **y** | **z** |
|---|---|---|---|---|---|---|
| Frontal lnf mid | <.001 | <.001 | 2891 | 54 | 32 | 35 |
| | | | | 46 | 50 | 11 |
| | | | | 52 | 50 | 25 |
| | | | | -13 | -12 | -2 |
| | | | | -13 | -1 | -4 |
| Frontal mid orb | <.001 | <.001 | 2728 | 1 | 66 | 3 |
| | | | | -4 | 72 | 11 |
| | | | | 2 | 57 | 0 |
| Frontal Sup medial | <.001 | <.001 | 1611 | 9 | 39 | 60 |

| | | | | | | |
|---|---|---|---|---|---|---|
| | | | | 18 | 52 | 57 |
| | | | | 6 | 55 | 58 |
| | <.001 | <.001 | 2296 | -8 | 65 | 52 |
| | | | | -4 | 50 | 59 |
| | | | | -6 | 58 | 47 |
| Frontal Inf Orb | <.001 | <.001 | 8695 | -44 | 44 | 5 |
| | | | | -53 | 43 | 26 |
| | | | | -52 | 37 | 20 |
| Frontal Inf Tri | 0.068 | 0.012 | 703 | 39 | 14 | 35 |
| | | | | 33 | 8 | 40 |
| | | | | 42 | 19 | 42 |
| Frontal Mid | 0.186 | 0.031 | 576 | -32 | 30 | 56 |
| | | | | -42 | 28 | 52 |
| Temporal Inf Lobe | <.001 | <.001 | 91628 | -44 | -35 | -26 |
| | | | | 58 | 18 | -3 |
| | | | | -44 | 42 | -3 |
| Lingual | <.001 | <.001 | 1757 | -16 | -36 | -7 |
| | | | | -11 | -25 | -10 |
| | | | | -24 | -32 | -8 |
| Cingulum Mid | 0.001 | <.001 | 1336 | -8 | -38 | 25 |
| | | | | -2 | -39 | 34 |

| | | | | | | |
|---|---|---|---|---|---|---|
| | | | | -14 | -33 | 27 |
| Amygdala | 0.005 | 0.012 | 1055 | -28 | 12 | -4 |
| | | | | -25 | 25 | 9 |
| | | | | -26 | 24 | -1 |
| Caudate | 0.037 | 0.012 | 782 | -10 | 28 | 19 |
| | | | | -22 | 18 | 20 |
| | | | | -17 | 36 | 20 |
| Cerebellum | 0.707 | 0.092 | 384 | -34 | -28 | -56 |
| Occipital- Primary visual Cortex | <.001 | <.001 | 1446 | 13 | -54 | -11 |
| **Small volume corrections emotional whole group** | | | | | | |
| Thalamus | 0.043 | 0.074 | 411 | -20 | -4 | 13 |

**Emotional younger adults study template (cl 383)**

| **Region** | **p(FWE-corr)** | **p(FDR-corr)** | **equivk** | **x** | **y** | **z** |
|---|---|---|---|---|---|---|
| Frontal sup | <.001 | <.001 | 12065 | -11 | 55 | 60 |
| | | | | -5 | 75 | 28 |
| | | | | -5 | 71 | 44 |
| Lingual | <.001 | <.001 | 8445 | -13 | -34 | -7 |
| | | | | -6 | -44 | -1 |
| | | | | 14 | -54 | -12 |

| | | | | | | |
|---|---|---|---|---|---|---|
| Occipital inferior | <.001 | <.001 | 6781 | 35 | -59 | -27 |
| | | | | 45 | -34 | -25 |
| | | | | 45 | -48 | -24 |
| Temporal superior | <.001 | <.001 | 1826 | 49 | 24 | -7 |
| | | | | 58 | 20 | -3 |
| | | | | 39 | 27 | -8 |
| Insula | <.001 | <.001 | 2294 | -24 | 25 | -1 |
| | | | | -29 | 10 | -7 |
| | | | | -32 | 21 | -14 |
| Cerebellum 9 | <.001 | <.001 | 1568 | -7 | -28 | -36 |
| | | | | 3 | -20 | -38 |
| | | | | 13 | -33 | -42 |
| Occipital inferior | <.001 | <.001 | 15913 | -36 | -61 | -30 |
| | | | | -32 | -66 | -24 |
| | | | | -14 | -45 | -31 |
| Temporal superior | <.001 | <.001 | 2471 | -60 | 2 | 1 |
| | | | | -54 | -1 | -8 |
| | | | | -47 | -4 | 1 |
| Frontal mid orb | <.001 | <.001 | 2487 | -28 | 37 | -19 |
| | | | | -44 | 42 | 4 |
| | | | | -45 | 42 | -4 |

| | | | | | | |
|---|---|---|---|---|---|---|
| Temporal mid | <.001 | <.001 | 1932 | -53 | -27 | 8 |
| | | | | -64 | -34 | 1 |
| | | | | -45 | -22 | 2 |
| Occipital inferior | <.001 | <.001 | 5162 | -23 | -73 | 0 |
| | | | | -13 | -77 | 5 |
| | | | | -32 | -70 | 11 |
| Temporal mid | <.001 | <.001 | 2443 | 57 | -5 | -3 |
| | | | | 50 | -11 | -1 |
| | | | | 49 | -21 | 0 |
| Insula | 0.512 | 0.03 | 433 | 22 | 24 | -4 |
| | | | | 33 | 20 | -2 |
| Caudate | 0.002 | <.001 | 1224 | 12 | 31 | 19 |
| | | | | 27 | 12 | 18 |
| | | | | 22 | 21 | 17 |
| Hippocampus | 0.019 | 0.001 | 867 | 28 | 11 | -8 |
| | | | | 20 | 8 | -11 |
| Frontal Inf orb | 0.688 | 0.048 | 383 | 28 | 28 | -15 |
| Temporal pole inf | | | | 31 | 38 | -24 |
| | | | | 25 | 31 | -23 |
| Caudate | 0.237 | 0.013 | 540 | -9 | 33 | 16 |
| | | | | -20 | 33 | 20 |

| | | | | | | |
|---|---|---|---|---|---|---|
| Cerebellum 9 | <.001 | <.001 | 3895 | 11 | -47 | -45 |
| Cerebellum Crus | | | | 19 | -52 | -39 |
| Cerebellum 8 | | | | 29 | -30 | -50 |
| Frontal Inferior TrI | | | | -52 | 46 | 21 |
| Cingulum posterior | 0.382 | 0.021 | 476 | 1 | -39 | 35 |
| Precuneus | | | | -6 | -44 | 39 |
| Temporal sup | 0.086 | 0.005 | 671 | 64 | -11 | 7 |
| | | | | 66 | -18 | 13 |
| | | | | 60 | -28 | 10 |
| Fusiform | | | | 36 | -16 | -25 |
| Temporal sup | 0.015 | 0.001 | 901 | -52 | 24 | 1 |
| Temporal pole sup | 0.187 | 0.011 | 571 | 45 | 36 | -18 |
| | | | | 42 | 29 | -21 |
| Frontal inf orb | 0.371 | 0.021 | 480 | 28 | 47 | -14 |
| | | | | 30 | 42 | -5 |
| | | | | 30 | 51 | -5 |
| Frontal mid orb | 0.092 | 0.005 | 661 | -22 | 25 | 47 |
| | | | | -27 | 23 | 55 |
| **Small volume corrections emotional younger adults study template** | | | | | | |
| Amygdala | 0.004 | 0.018 | 572 | -27 | 11 | -6 |
| Hippocampus | 0.01 | 0.025 | 582 | 20 | 8 | -11 |

| **Emotional older adults study template (cl 417 )** | | | | | | |
|---|---|---|---|---|---|---|
| **Region** | **p(FWE-corr)** | **p(FDR-corr)** | **equivk** | **x** | **y** | **z** |
| Frontal Inf Tri | 0.005 | 0.001 | 940 | -47 | 51 | 16 |
| | | | | -39 | 51 | 17 |
| Temporal Mid Lobe | 0.001 | <.001 | 1223 | -56 | -23 | -4 |
| | | | | -62 | -31 | -2 |
| | | | | -64 | -22 | -2 |
| Temporal Inf | 0.003 | 0.001 | 998 | -48 | -30 | -25 |
| | | | | -44 | -22 | -21 |
| | | | | -51 | -40 | -24 |
| Temporal Sup | <.001 | <.001 | 1873 | -56 | -30 | 14 |
| | | | | -48 | -33 | 17 |
| | | | | -58 | -40 | 9 |
| Precuneus | 0.348 | 0.035 | 448 | -10 | -38 | 22 |
| Occipital Inf | 0.015 | 0.002 | 811 | 37 | -66 | -15 |
| | | | | 41 | -59 | -22 |
| | | | | 32 | -70 | -8 |
| | 0.086 | 0.022 | 610 | -40 | -65 | -10 |
| | | | | -41 | -63 | -18 |
| | | | | -36 | -69 | -1 |

| | | | | | | |
|---|---|---|---|---|---|---|
| Occipital Mid | 0.096 | 0.022 | 598 | -50 | -49 | 24 |
| | | | | -42 | -58 | 22 |
| | | | | -46 | -48 | 17 |
| Cerebellum | <.001 | <.001 | 5522 | 20 | -45 | -49 |
| | | | | -20 | -43 | -45 |
| | | | | 19 | -40 | -41 |
| **Small Volume Corrections emotional older adults study template** | | | | | | |
| No small structures significant (No SVC) | | | | | | |
| **Emotional MCI study template (cl 820)** | | | | | | |
| **Region** | **p(FWE-corr)** | **p(FDR-corr)** | **equivk** | **x** | **y** | **z** |
| Temporal Mid | 0.017 | 0.01 | 844 | -56 | -42 | 5 |
| | | | | -53 | -51 | 2 |
| Occipital Inf | 0.021 | 0.01 | 820 | 49 | -42 | -20 |
| | | | | 43 | -45 | -25 |
| | | | | 54 | -38 | -12 |
| **Small volume corrections emotional MCI study template** | | | | | | |
| No small structures significant (No SVC) | | | | | | |
| **Emotional: additional results** | | | | | | |
| **Emotional younger adults > older adults study template (cl 745)** | | | | | | |
| No suprathreshold clusters | | | | | | |

**Emotional older adults > younger adults study template**

FDRC Inf.

No small structures significant (No SVC)

**Emotional older adults > MCI study template**

FDRC Inf.

No small structures significant (No SVC)

**Emotional MCI > older adults study template**

FDRC Inf.

No small structures significant (No SVC)

**Table 3. Significant activations Remembered Images Larger than not Remembered Images (memory)**

**Memory whole group study template**

| | | | | | | |
|---|---|---|---|---|---|---|
| FDRC inf | | | | | | |
| **Small volume corrections memory whole group** | | | | | | |
| Putamen | 0.005 | 0.008 | 665 | 19 | 29 | 6 |
| Parahippocampus | 0.121 | 0.136 | 206 | -32 | 1 | -17 |
| | 0.102 | 0.136 | 225 | 26 | -16 | -12 |

**Memory younger adults study template (cl 482)**

| **Region** | **p(FWE-corr)** | **p(FDR-corr)** | **equivk** | **x** | **y** | **z** |
|---|---|---|---|---|---|---|
| Precentral | <.001 | <.001 | 2416 | 35 | -6 | 41 |
| | | | | 39 | -6 | 51 |
| | | | | 46 | -6 | 42 |
| Parahippocampus | 0.165 | 0.018 | 558 | -33 | 0 | -15 |
| | | | | -36 | -11 | -10 |
| Parahippocampus | <.001 | <.001 | 2123 | 36 | -13 | -9 |
| | | | | 25 | -22 | -10 |
| | | | | 31 | -11 | -16 |
| Lingual | <.001 | <.001 | 2338 | -29 | -25 | -16 |
| | | | | -10 | -25 | -23 |
| | | | | -25 | -31 | -20 |
| Cuneus | 0.048 | 0.007 | 708 | 17 | -54 | 34 |
| | | | | 47 | -58 | 4 |
| | | | | 33 | -62 | -5 |
| Occipital mid | <.001 | <.001 | 1776 | -21 | -72 | -5 |
| | | | | -35 | -70 | -4 |
| | | | | -17 | -68 | -13 |
| Occipital mid R | 0.306 | 0.033 | 482 | 38 | -61 | 2 |
| | | | | 47 | -58 | 4 |

| | | | | | | |
|---|---|---|---|---|---|---|
| | | | | 33 | -62 | -5 |
| **Small volume corrections memory younger adults study template** | | | | | | |
| Putamen | 0.111 | 0.151 | 272 | 18 | 25 | 5 |
| **Memory older adults study template (cl 627)** | | | | | | |
| **Region** | **p(FWE-corr)** | **p(FDR-corr)** | **equivk** | **x** | **y** | **z** |
| Putamen | 0.058 | 0.027 | 627 | 24 | 22 | 3 |
| **Small volume corrections memory older adults** | | | | | | |
| LC | 0.082 | 0.936 | 4 | -6 | -8 | -27 |
| Putamen | 0.047 | 0.041 | 335 | -23 | 32 | 8 |
| | 0.002 | 0.004 | 725 | 24 | 22 | 3 |
| **Memory MCI study template** | | | | | | |
| Inf | | | | | | |
| No SVC | | | | | | |
| **Memory: additional results** | | | | | | |
| **Memory older adults > MCI study template** | | | | | | |
| Inf | | | | | | |
| **Small volume corrections memory older adults > MCI study template** | | | | | | |
| LC | 0.081 | 0.936 | 4 | -6 | -8 | -27 |
| **MCI > Older adults** | | | | | | |
| Inf | | | | | | |

No SVC

| **Memory younger adults > older adults study template (cl 637)** | | | | | | |
|---|---|---|---|---|---|---|
| **Region** | **p(FWE-corr)** | **p(FDR-corr)** | **equivk** | **x** | **y** | **z** |
| Cerebellum 7b | <.001 | 0.01 | 1360 | -44 | -14 | -44 |
| | | | | -43 | -28 | -54 |
| | | | | -38 | -17 | -57 |
| Precuneus | 0.002 | 0.01 | 1130 | 11 | -58 | 34 |
| | | | | 20 | -52 | 39 |
| | | | | 9 | -64 | 23 |
| Precentral | 0.096 | 0.015 | 644 | 35 | -1 | 54 |
| | | | | 37 | -5 | 45 |
| | | | | 43 | -4 | 53 |

| **Memory younger adults > MCI study template (cl 716)** | | | | | | |
|---|---|---|---|---|---|---|
| **Region** | **p(FWE-corr)** | **p(FDR-corr)** | **equivk** | **x** | **y** | **z** |
| Cuneus | 0.055 | 0.015 | 716 | 17 | -53 | 35 |
| | | | | 27 | -51 | 35 |
| Fusiform | 0.014 | 0.006 | 889 | -29 | -25 | -16 |
| | | | | -21 | -25 | -13 |
| | | | | -31 | -19 | -11 |

| | | | | | | |
|---|---|---|---|---|---|---|
| Precentral | 0.002 | 0.002 | 1156 | 37 | -5 | 47 |
| | | | | 46 | -2 | 55 |
| | | | | 34 | -11 | 39 |

**Small volume corrections memory younger adults > older adults study template**

No SVC

**Memory older adults > younger adults study template**

Inf

.01 Inf - no SVC

**Memory MCI > younger adults study template**

Inf
